# Evolutionary and functional genomics reveal that *Ralstonia* wilt pathogens actively deploy antimicrobial warfare while leveraging physiological adaptations during plant infection

**DOI:** 10.64898/2026.03.20.712958

**Authors:** Nathalie Aoun, Stratton J. Georgoulis, Adam M. Deutschbauer, Tiffany M. Lowe-Power

## Abstract

The xylem pathogens in the *Ralstonia solanacearum* species complex (RSSC) cause wilt diseases that threaten global food security. These diverse pathogens are highly adapted to the *in planta* environment where they proliferate and cause rapid, aggressive diseases. To understand the genetic underpinnings of these pathogens’ *in planta* fitness, we performed forward genetic screens in three RSSC species, using high-throughput random barcode transposon sequencing (RB-TnSeq). We quantified the competitive fitness of hundreds of thousands of mutants during stem colonization of susceptible tomato plants. We compared this *in planta* forward genetic screen with a prior forward genetic screen performed in a reductionist, plant-like environment: *ex vivo* xylem sap harvest from healthy tomato plants. The comparative genetic screens revealed both conserved and lineage-specific orthologs that enable RSSC’s pathogenic success. Consistently, these diverse pathogens navigated the opportunities and stressors *in planta* by maintaining a resilient cell envelope barrier, complementing the nutritional availability in xylem sap, fine-tuning their utilization of the host-manipulating type III secretion system, and dynamically regulating their gene expression through a suite of environment sensing and transcriptional regulatory proteins. Surprisingly, the strain-specific fitness factors shed light on ecological interactions beyond pathogen dominance of their susceptible host, even during the single-strain infections, the pathogens wielded arsenals of anti-microbial weapons. Specifically during growth *in planta,* each RSSC strain required a unique repertoire of type VI secretion system immunity proteins that provide self-protection to their own toxins. We contextualized the *in planta* fitness factors through evolutionary genomic comparisons of the RSSC wilt pathogens to their non-pathogenic neighbors. Together, these analyses reveal how conserved and lineage-specific fitness determinants have evolved to support pathogenic success in the plant vascular niche.

## Introduction

Bacterial wilt caused by the soil-borne *Ralstonia solanacearum* species complex (RSSC) pathogens is a devastating vascular disease affecting a wide range of plant hosts [1,2]. RSSC are soil-borne pathogens that locate host roots using chemotaxis and flagellar motility [3,4]. RSSC pathogens invade the root through structurally weak tissue, such as the lateral root emergences [5,6]. The bacterial pathogens navigate to the vascular cortex where they invade the xylem. In the xylem, the pathogens proliferate and rapidly disseminate vessel-to-vessel to systemically colonize and ultimately clog vessels, leading to wilt symptoms [7]. Throughout infection, the pathogens shed from the roots to return to soil [8]. Extensive work over the past two decades has revealed the complexity of RSSC pathogenicity using system-level approaches although most mechanistic studies focus on a single strain background [9–18].

Plants transport water from roots to shoots using pressure differentials across the xylem vessel network [19]. Although the xylem vessels are a physiologically dead tissue, the metabolite and protein components of xylem sap are controlled by the adjacent living parenchyma cells. To spread vessel-to-vessel, RSSC secrete a suite of cellulases and pectinases to degrade the pit membranes between adjoining vessels [20,21]. Additionally, these cell-wall-degrading enzymes break down the pit membranes that separate vessels from the living parenchyma tissue, allowing the pathogens to spill out into the stem apoplast [22].

One approach to understanding how RSSC thrive during infection is to use reductionist experiments where the bacteria are grown in *ex vivo* xylem sap. For example, we previously conducted a high-throughput forward genetic screen to identify genes that contribute to pathogen growth in *ex vivo* xylem sap from susceptible tomato plants [23]. Although reductionist approaches are valuable, RSSC pathogens experience an *in planta* environment that is physically, chemically, and biologically different from the *in vitro* environment [24]. These environmental transformations occur through direct actions by the pathogen, indirect effects caused by pathogen damage to the host, and host-mediated changes following immune perception of the pathogen.

Many mechanistic pathogenesis studies have identified traits that promote *in planta* success of RSSC wilt pathogens. Due to time and cost constraints, detailed mechanistic studies typically use a single model strain to explain pathogenesis. However, within the RSSC, there is considerable genetic, phenotypic, and ecological diversity [25–27]. The RSSC pathogens are a monophyletic clade of three wilt-pathogenic species within the *Ralstonia* genus: *R. solanacearum* (Rsol), *R. pseudosolanacearum* (Rpseu), and *R. syzygii* (Rsyz) [26]. Across the species diversity, these pathogens have complex patterns of generalist host range that vary dramatically between strains and lineages [1,2,28]. Nevertheless, most known lineages convergently evolved in ways that have enabled them to share pathogenicity to Solanaceae family plants, including the socioeconomically important crop tomato [28].

Comparative evolutionary approaches between pathogens and their evolutionary neighbors can differentiate ancestral genes and traits associated with the emergence of pathogenesis from the genes and traits gained later in evolution [29]. In addition to the RSSC, the *Ralstonia* genus contains a second major clade of dozens of species with less-well characterized ecological roles [30,31]. These evolutionary neighbors have been isolated from habitats that overlap with RSSC pathogens’ habitats (freshwater, soil, and plant rhizosphere) and are unique to the neighbors (industrial water systems and opportunistic infections of humans). Because these neighboring species share ancestry and partially overlapping habitats with RSSC pathogens, they provide a powerful comparative framework for distinguishing traits associated specifically with plant pathogenesis [32].

In this study, we performed several genome-wide forward genetic screens in three RSSC strains, each capable of causing bacterial wilt disease of tomato. Using a barcoded mutant approach, we competitively grew hundreds of thousands of mutants in stems of susceptible tomato plants following direct inoculation into the stem xylem. To contextualize the identified fitness factors, we compared and contrasted this study’s *in planta* competitive fitness results with our previously published genetic screens in *ex vivo* xylem sap from healthy tomato plants of the same cultivar [23]. Genomic comparisons provided insight into the evolutionary history of these fitness factors. Together, this study reveals a comprehensive fitness landscape that reflects both the conserved and diverse strategies of these heterogeneous, globally important plant pathogens.

## Materials and Methods

### Bacterial Strains, Mutant Libraries, and Growth Conditions

This study uses wildtype strains and barcoded mariner transposon mutant libraries (“RB-TnSeq libraries”) created in the three RSSC strain backgrounds *R. pseudosolanacearum* GMI1000, *R. solanacearum* IBSBF1503, and *R. syzygii* PSI07. The construction and properties of the RB-TnSeq libraries are described previously [23]. Bacteria were routinely grown in CPG rich medium, containing per 1 liter: 1 g casamino acids, 10 g Bacto peptone, 5 g glucose, and 1 g yeast extract. The RB-TnSeq libraries were cultured with “half-strength” kanamycin at 12.5 μg/mL in case mutants had increased sensitivity to the antibiotic.

### Preparation of RB-TnSeq Libraries for Competitive Fitness Screens

To prepare inoculum for RB-TnSeq experiments, a 1 mL cryostock aliquot of the barcoded mutant libraries was thawed on ice for 30 min, inoculated into 100 mL CPG with 12.5 μg/mL kanamycin in a 250 mL flask, and incubated at 28 °C with shaking. Once the culture reached the mid-log phase (OD_600_ between 0.4 and 0.7), bacterial cells were pelleted. In experiment 1 (described below), pellets were resuspended in the remaining culture supernatant. In experiments 2 and 3 (described below), pellets were washed and resuspended in 1 mL of MilliQ ultrapure water. The cell density of the washed, revived mutant libraries was adjusted to 5x10^10^ CFU/mL to yield a target inoculum of 10^8^ CFU per plant. Duplicates of 10^9^ cells inoculum (20 μl) were saved and frozen at -20 °C as time_0_ controls. To directly inoculate plant stems, we used a sharp blade to remove the oldest petiole above the cotyledons and inoculated a 2 μl droplet onto the stump; within minutes the negative pressure in the xylem network pulled the droplet into the xylem. To directly inoculate plant stems, we used a sharp blade to remove the oldest petiole above the cotyledons and applied a 2 µl droplet of inoculum onto the exposed stump. Within minutes, negative pressure in the xylem network pulled the droplet into the vascular tissue. Empirical measurements indicated that most inoculated cells entered the stem during these cut-petiole inoculations; by 1 hour after inoculation, approximately half of the original inoculum remained viable following a 2-minute surface treatment with 10% bleach, consistent with internalization. Harvesting of the mutant populations is described below.

Over four years of experiments, the recovery time for cryopreserved IBSBF1503 and PSI07 libraries increased from 19 to 24 hr., suggesting high viability. In contrast, the recovery time for the cryopreserved GMI1000 library gradually increased from 18 to 36 hr, suggesting that half of the cryopreserved populations lost viability. Despite the extended recovery time, all time_0_ samples contained high barcode diversity and abundance, and dilution plating demonstrated that atypical colony morphologies (including quorum sensing mutants) remained rare.

### Experimental Design for RB-TnSeq Screens in Stems of Susceptible Tomato Plants

We carried out three experimental designs to identify which genes promote or hinder RSSC growth in stems of tomato *cv.* Moneymaker plants (Fig. S1). In all experiments, the inoculation, as described above, was consistent. Additionally, to sample the total stem, the stem was cut near the soil line, and the petioles were cut just below the first leaflet. Otherwise, the experiments varied by plant growth environment, plant size, and plant tissues sampled (pooled stems and pooled roots, pooled stems only, or stems from individual plants). The timing of bacterial harvest varied from 3 to 5 days following inoculation, as described below.

**Experiment 1:** N=3 biological replicates, each consisting of 20 individually inoculated plants that were pooled together during harvest. The total stem and root system were both harvested, yielding “proximal site” (stem after cut-petiole inoculation) and “distal site” (roots after cut-petiole inoculation). Although the bacteria were harvested from roots in the distal condition, this dataset should not be mistaken for a screen identifying genes required for bacterial root invasion from the soil.

Tomato seeds were sown in potting mix and grown in the Oxford Tract Greenhouse in Berkeley, CA. After 14 days, plants were transplanted into 4-inch pots. Once the plants reached the 3-4 leaf stage (20-35 days due to seasonal fluctuations in greenhouse conditions), plants were transferred to a Conviron growth chamber (“Growth Chamber 1”) maintained at 28°C and a 12 hr day/night cycle. Plants were watered with an in-house nutrient solution while in the greenhouse and with tap water while in the growth chamber. After plants had acclimated to the growth chamber for 4 days, the RB-TnSeq inoculations were carried out as described above.

At 3 days post-inoculation, total plant stems and roots were harvested. Plants were removed from their pots, and bulk soil was removed by gentle shaking. The rhizosphere soil was removed by gently agitating the roots in water. The roots were briefly surface sterilized by gentle agitation in 10% bleach for 30 s. The remaining bleach was removed by gently agitating the roots in three sequential containers of water for 30 s. The roots were blotted, dried, weighed, and homogenized in 100 mL of MilliQ ultrapure water in a Magic Bullet® blender.

Separately, the total stems were harvested by removing leaves, coarsely chopping the stem, and then homogenizing like the roots. Blender canisters were thoroughly sterilized between samples by soaking in 10% bleach, followed by rinsing. To reduce plant debris, the homogenate was filtered sequentially through coffee filters, filters with 20 μm pores (Whatman™ 1004-070), and 6 μm pores (Whatman™ 1003-070). Pellets containing bacteria were collected from the filtrate by centrifugation (16,000 rcf) and frozen at -20°C until DNA extraction.

**Experiment 2:** Experiment 2 was an experimental repeat of Experiment 1 but using a different growth chamber. Here, only the proximal sample of total stem were harvested. There were N=3 biological replicates that each consisted of 18 individually inoculated and pooled plants. Tomato seeds were sown in potting mix and grown in a growth chamber in the Controlled Environment Facility at Davis, CA maintained at 28°C and a 12-hr. day/night cycle.

After 14 days, seedlings were transplanted into 3-inch pots. Plants were watered with an in-house nutrient solution at all stages. RB-TnSeq inoculations and sampling were carried out as described above with minor modifications due to the timing of wilting symptoms. After 3, 4, and 5 days after inoculation with the GMI1000, IBSBF1503, and PSI07 libraries, respectively.

**Experiment 3:** This experiment had N=5 biological replicates, and each replicate was the stem from an individual plant. Plants were grown and inoculated as described for Experiment 2. Stems from individual plants were collected at 4 days (GMI1000 and IBSBF1503) at 5 days (PSI07).

### Barcode Sequencing and Fitness Scores For RB-TnSeq Screens

DNA was extracted from time_0_ and pelleted cells from homogenized stems using the Qiagen DNeasy Powerlyser PowerSoil Kit. PCR was used to amplify the DNA barcodes and the barcodes were sequenced with Illumina HiSeq4000 at QB3 Berkeley Genomics Center with 96 samples multiplexed (50-bp reads, single end), as described previously [33]. Two technical replicates were sequenced per biological replicate for experiment 1 while one technical replicate was sequenced per biological replicate for experiments 2 and 3. For each experimental condition, fitness scores for each gene were calculated as described previously [33]. A fitness score corresponds to the log_2_ ratio between barcoded mutants’ abundance after competitive growth in the condition, divided by the same strains’ abundance in the time_0_ sample. Fitness scores are integrated into the online Fitness Browser (fit.genomics.lbl.gov) for samples that passed stringent quality control metrics; full quality control metrics are present in Tables S1-S3.

### Identification of Syntenic Orthologs in GMI1000, IBSBF1503, and PSI07

Syntenic orthologs in the three strains were identified by manual curation of pangenome and synteny results. The RefSeq assemblies for GMI1000, PSI07, and IBSBF1503 were compared using the KBase “Compute Pangenome” pipeline [34]. Briefly, this pipeline uses a protein k-mer based similarity approach to cluster proteins into putatively homologous families when they share at least five unique 8-mers (after filtering out any 8-mer occurring more than 5 times in a single genome) and then rescans and rescores preliminary clusters to split overly heterogenous protein families. Clinker was used to developed interactive, synteny plots across eleven chromosome regions and five megaplasmid regions spanning the bipartite genomes [35]; the interactive synteny plots color-code homologous genes and link them with visualizations that convey pairwise global amino acid identity values. The .gbk input files for clinker were extracted from the “graphics” page of the NCBI nucleotide database for each RefSeq accession. The synteny plots were manually inspected to curate a final table of syntenic orthologs and singletons.

### Identification of shared and strain-specific *in planta* fitness factors

Fitness scores were plotted as boxplots using ‘ggplot2’ [36] in R. Correlated plots among RB-TnSeq conditions were generated using ‘cor’ and plotted using ‘ggplot2’ in R (Figs. S2-S7). To identify genes that contribute to any of the RSSC strains’ *in planta* growth, we focused on pooled-stem samples from experiments 1 and 2. Biological replicates were assessed for consistency based on pairwise Pearson correlation coefficients; one replicate from the GMI1000 and one replicate from the IBSBF1503 experiments had low correlation were excluded from downstream analyses. For each retained replicate, we selected the 500 genes with the highest fitness scores and the 500 genes with the lowest fitness scores. Genes were retained only if they appeared in ≥4 replicates for *R. pseudosolanacearum* GMI1000 and *R. syzygii* PSI07, or ≥5 replicates for *R. solanacearum* IBSBF1503. The data on syntenic orthologs were aggregated to generate a unified list of candidate fitness factors, and we retained genes for which at least one ortholog had an average absolute fitness score ≥ 0.5.

### Assigning Functional Categories to Genes

Preliminary functional categories were assigned to fitness factors using the NCBI Cluster of Orthologous Group (COG) database [37,38]. We manually checked the automated assignments and tailored categories based on expert opinion and PaperBlast results. For example, we moved *fca* and the *eps* genes from “Function unknown” to “Phenolic transport and metabolism” [39] and “EPS-I exopolysaccharide” [40], respectively.

### Construction of Phylogenetic Species Trees

We used KBase to create a phylogenetic trees by using the pipeline Insert Set of Genomes into a SpeciesTree-v.2.2.0, which identifies 49 COGs that are universally conserved in single-copy across bacteria, concatenates the sequences, builds an MSA, and creates an approximate maximum likelihood tree with FastTree2 [34,41]. The Newick file output was downloaded and imported into iTOL [42] for visualization, and phylogenetic trees were polished using Affinity Designer.

### Analysis of Evolutionary Conservation of Stem Fitness Factors

To explore the conservation of fitness factors, a pangenome analysis was performed on a phylogenetically balanced set of representative RSSC phytopathogen (n=24) and environmental *Ralstonia* spp. (n=29) genomes. Balanced genomes were selected from a 49-gene phylogenetic tree of all publicly available *Ralstonia* genomes as of 2024 [1]. These genomes had < 2.03% contamination and were ≥99.48% complete based on the CheckM markersets for the Burkholderiaceae family [43]. Within KBase, mOTUpan v0.3.2 was run for 1 iteration (--max_iter 1) using cut-offs for 75% identity and 80% coverage (MMseqs2 parameters: easy-cluster --min-seq-id 0.75 -c 0.80) [44]. The percent identity cut-off of 75% was selected based on empirical results from running the analysis with 60%, 70%, 75%, and 80% identity cut-offs.

For a subset of genes, conservation was explored the BLASTp searches against high-quality, publicly available RSSC genomes (n=621; available at https://narrative.kbase.us/narrative/189849) and environmental *Ralstonia* spp. genomes (n=150; available at https://narrative.kbase.us/narrative/189428). To filter out low-quality genomes, we ran CheckM-*v1.0.18* [43] to estimate the completeness and contamination of all genomes. Genomes with completeness lower than 99.82% and contamination greater than 0.96% were discarded. Genomospecies designations were applied across all strains using a 95% ANI threshold. Unnamed species were assigned Genome Taxonomy Database (GTDB) placeholder designations [45]. We queried protein sequences fitness factors against the RSSC and environmental *Ralstonia* genomes using BLASTp-*v2.13.0* in KBase. The results of the BLASTp hits were filtered to identify the highest identity hit with at least 60% query coverage. The results were imported into iTOL to visualize the phylogenetic distributions. We routinely performed synteny analysis using Clinker to confirm that hits were likely to be ortholog [35].

### Disease progress assays

Cut-petiole inoculations of tomato *cv.* Moneymaker plants with the three RSSC wild-type strains were performed in three biological replicates with 18 plants per strain. When plants were 21 days-old, they were cut-petiole inoculated ∼1000 CFU per plant. Symptoms were measured daily for 14 days using a standard 0-4 disease index scale where 0: no symptoms; 1: up to 25% leaflets wilted; 2: up to 50% of leaflets wilted; 3: up to 75% of leaflets wilted; 4: up to 100% of leaflets wilted [46].

## Results

### High-throughput genetic screens identified conserved and strain-specific *in planta* fitness factors for three wilt pathogenic RSSC species representatives

To investigate genetic factors contributing to the *in planta* growth of wilt pathogenic RSSC, we performed a high-throughput genetic screen with barcoded Tn-insertion mutants (RB-TnSeq) [33]. To capture the genetic diversity of fitness traits across phylogenetic lineages, we performed RB-TnSeq screens in the genetic backgrounds of three tomato-pathogenic strains that represent the three species: *R. pseudosolanacearum* (Rpseu) GMI1000, *R. solanacearum* (Rsol) IBSBF1503, and *R. syzygii* (Rsyz) PSI07 (Fig. 1a-b and Fig. S1).

**Figure 1.**
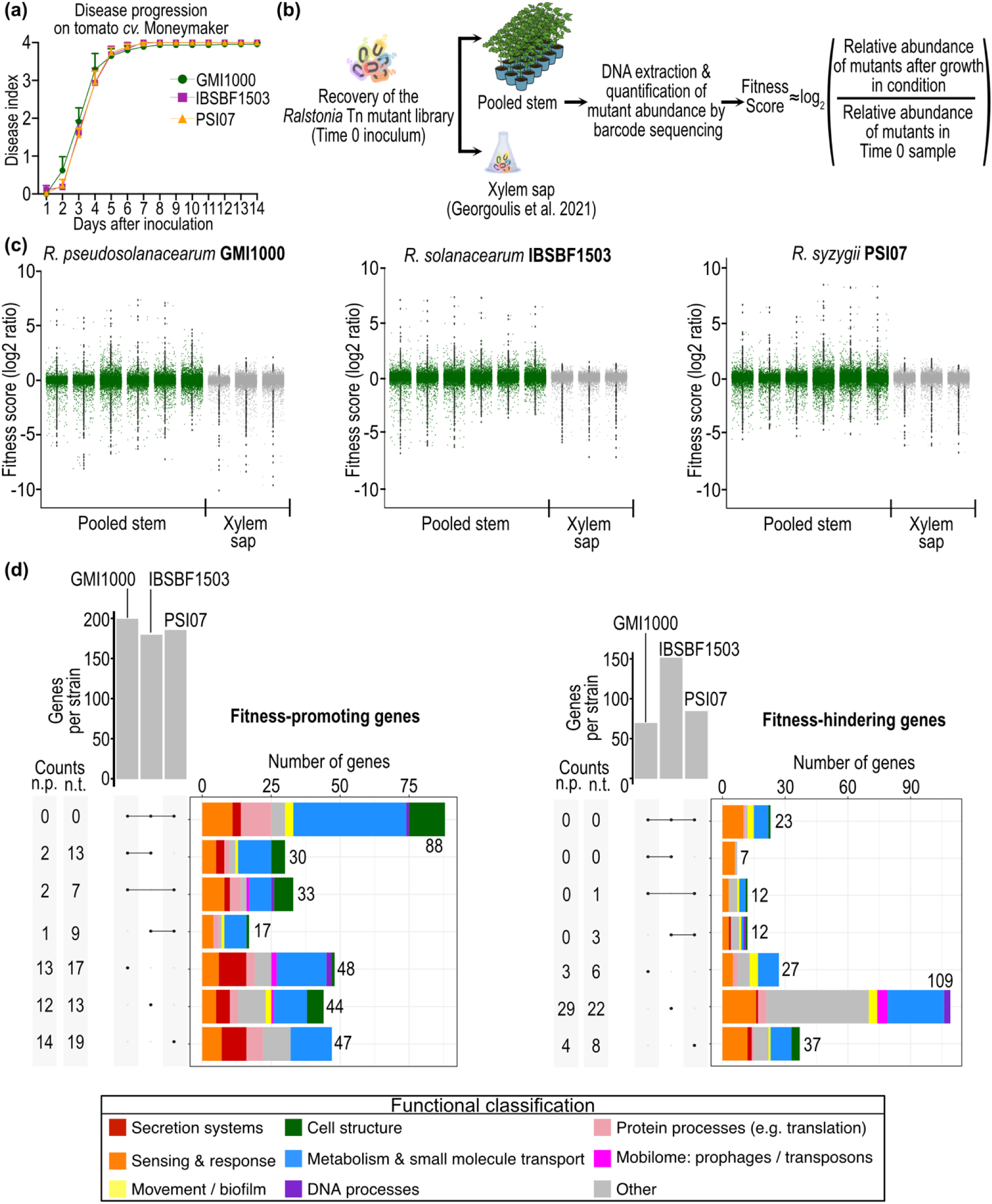
Genome-wide screen for factors that promote or hinder growth of diverse plant pathogenic RSSC in susceptible tomato *cv.* Moneymaker plants. (**a**) Disease progress following cut-petiole inoculation of tomato *cv.* Moneymaker plants with three RSSC species representatives: Rpseu GMI1000, Rsol IBSBF1503, and Rsyz PSI07. Symptoms were monitored daily for 14 days using a 0-4 disease index scale: 0, no wilted leaflets (0%); 1, >0–25% wilted leaflets; 2, >25–50% wilted leaflets; 3, >50–75% wilted leaflets; and 4, >75–100% wilted leaflets. The experiment was performed in three biological replicates with n=18 plants per replicate. Symbols represent the mean, and error bars indicate the standard error of mean (SEM). (**b**) Overview of the experimental design for high-throughput genetic screens using random barcoded transposon mutant sequencing (RB-TnSeq). (**c**) Distribution of RB-TnSeq fitness scores for the three RSSC species representatives in stems for susceptible tomato (n=6 experimental replicates) vs. in *ex vivo* xylem sap from health tomato plants (n=3 experimental replicates from Georgoulis et al., 2021. (**d**) Comparison of genes that promoted or hindered *in planta* growth in at least one RSSC species representative background (threshold |Fitness Score|<0.5). UpSet plots display the total number of fitness-impacting genes per strain (vertical bar charts) and the number of genes with fitness effects among all three, two, or only one RSSC species representatives (horizontal stacked bar charts colored based on functional classification of the fitness factors). Columns to the left of each UpSet plot indicate genes excluded from the intersection analysis due to the strain naturally lacking an ortholog (“n.p.”, not present), or due to missing fitness data caused by stochastic absence of the corresponding mutant in the RB-TnSeq library (“n.t.”, not tested). UpSet plots were generated using “UpsetR” and “ggplot2” in R. Figures were refined in Affinity Designer.

We inoculated the mutant libraries directly into stem xylem of susceptible tomato plants (*cv.* Moneymaker) via a cut-petiole inoculation. For six biological replicates per strain, the mutant populations were harvested from pooled stems from 18-20 tomato plants after several days when many plants displayed wilt symptoms of this rapid disease. Based on dilution plating, the bacterial populations doubled between 4 and 13 times, generally exceeding 1.2x10^13^ cfu/plant total stem. Mutant fitness scores were calculated as a log_2_ ratio of the relative abundance of sequenced barcodes in the populations after *in planta* growth vs. their abundance in the starting inoculum (Tables S1-S3). Additionally for the first three replicates of the pooled stem experimental design, we obtained RB-TnSeq fitness profiles obtained from mutants recovered from roots-after-stem inoculation (distal site) in addition to the stem-after-stem inoculation (proximal site). The results from this distal site were largely the same as the results from the proximal site, so we focused subsequent analyses on the stem dataset.

Fitness scores of many replicates were highly correlated, especially in the genetic background of Rpseu-GMI1000 and Rsol-IBSBF1503 (Figs. S2-S4). However, one of the six Rpseu-GMI1000 and one of six Rsyz-PSI07 replicates were removed from further analysis because the outlier replicates had both a low correlation with other replicates and/or a narrow distribution of fitness scores (Figs. 1c, S1, and S4); all six replicates of Rsol-IBSBF1503 were retained. Across the retained biological replicates, the average genic fitness scores ranged from -5.7 to +6.1 (Rpseu-GMI1000), -4.9 to +6.8 (Rsol-IBSBF1503), and -4.8 to +6.8 (Rsyz-PSI07) (Fig. 1c and Tables S1-S3). A negative fitness score indicates that the mutant was depleted *in planta* relative to the total population, suggesting that the corresponding gene promotes pathogen growth *in planta*. Conversely, positive fitness scores correspond to mutants that were enriched, suggesting that the corresponding gene hinders RSSC growth *in planta*. Thus, the RB-TnSeq screens identified mutants that grew up to 50-fold worse and 158-fold better than expected for a wildtype cell.

We explored whether the RB-TnSeq screens yield consistent results when the mutants were harvested from individual plants, compared to the pooled plant samples. Fitness scores from the individually harvested plants spanned a wider range: -10.0 to +9.9 in Rpseu-GMI1000, -6.1 to +7.0 in Rsol-IBSBF1503, and -5.5 to +9.9 in Rsyz-PSI07. Among the individual plant replicates, the samples were well-correlated in Rpseu-GMI1000 (*r* = 0.55-0.76) and Rsol-IBSBF1503 (*r* = 0.36-0.57), but the Rsyz-PSI07 replicates were poorly correlated with each other (*r* = 0.18-0.27) (Figs. S5-S7). When comparing the two experimental designs, the pooled and individual plant replicates were only well-correlated to each other in Rsol-IBSBF1503 (*r* = 0.29-0.58) (Fig. S6). Because the pooled samples yielded more consistent results (Rpseu-GMI1000: *r* =0.48-0.86, Rsol-IBSBF1503: *r* =0.47-0.85, and Rsyz-PSI07: *r* =0.25-0.75), we focused further analysis on the pooled plant replicates.

After assessing replicate consistency, we next sought to systematically identify genes that influence *in planta* fitness in the pooled samples across strains. Because differences in population growth among replicates affect fitness score magnitudes, defining thresholds for biological significance presents a persistent analytical challenge in TnSeq studies. To address this issue and to ensure comprehensive identification of fitness factors across strains, we implemented a holistic analytical framework before examining strain-to-strain variation (Fig. S8). We selected the genes with either the 500 lowest or the 500 highest fitness scores in each retained biological replicate (Fig. S8). We then filtered this list for genes that consistently appeared in most replicates (≥4 for Rpseu-GMI1000 and Rsyz-PSI07; and ≥5 for Rsol-IBSBF1503). We identified syntenic orthologs, allowing aggregation of the top hits from all three strain backgrounds. To focus on biologically meaningful effects, we filtered the dataset to genes in which at least one ortholog had an average |Fitness Score| ≥ 0.5 (Table S4).

Overall, more RSSC mutants were depleted than enriched *in planta* (N=307 and N=227, respectively) (Fig. 1c and Table S4). We assigned functional categories using a combination of Clusters of Orthologous Groups (COG) analysis [37,38], PaperBLAST [47], and manual curation of RSSC-specific functions (Figs. 1d-3 and Table S4). Diverse biological functions influenced RSSC growth, including protein secretion system-related genes, environmental sensing and response pathways, movement and biofilm factors, cell envelope and cell structure, metabolism and transporters, DNA repair, translation and other protein processes, mobilome and mobile genetic element defenses, and other or unknown functions.

**Figure 2.**
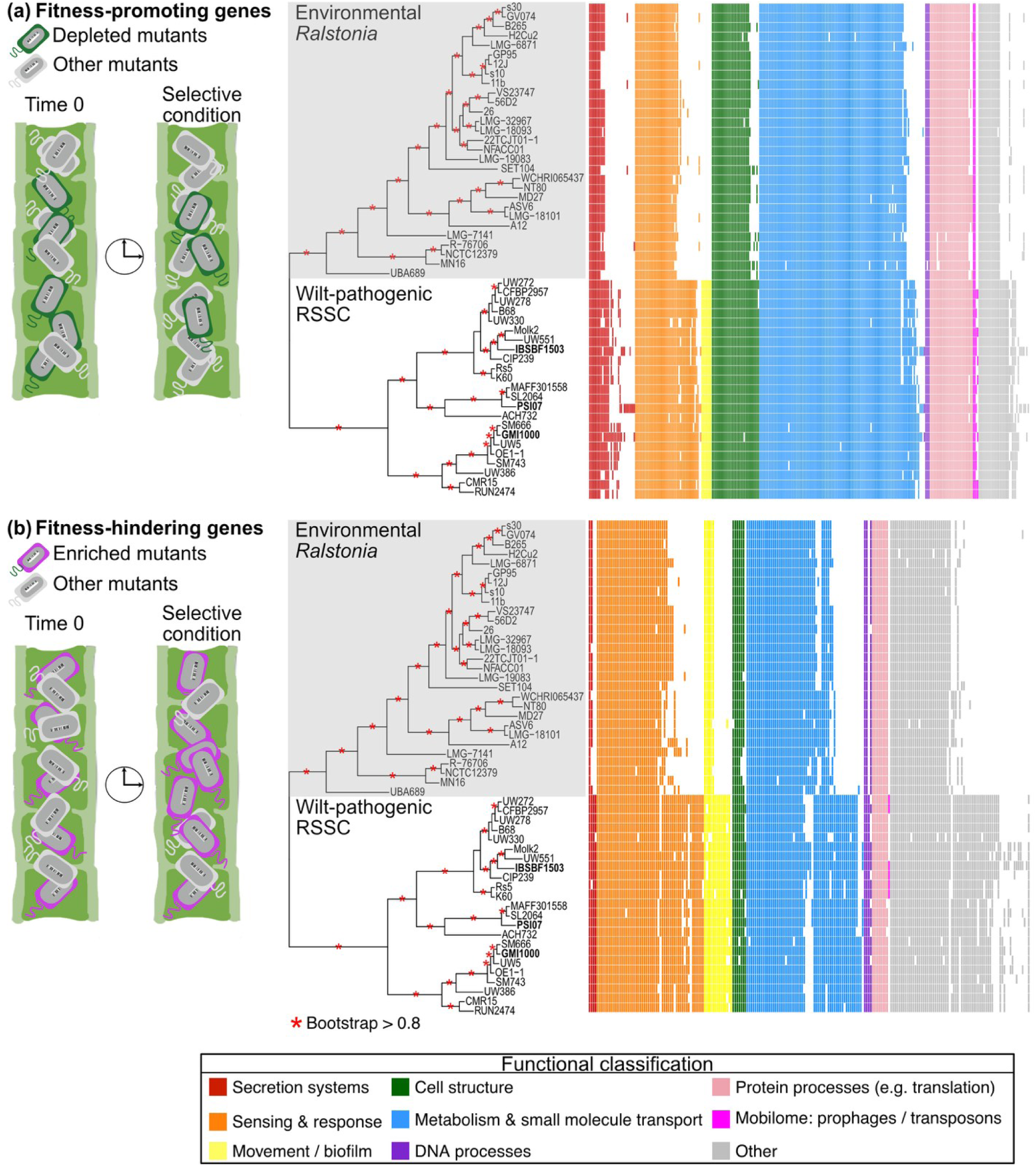
Genetic conservation of *in planta* fitness factors highlights RSSC adaptations to the wilt pathogenic lifestyle. Pangenome analysis of the *Ralstonia* genus was performed to investigate the genetic conservation of RSSC *in planta* fitness factors that (**a**) promote or (**b**) hinder *in planta* growth. Orthologs were identified using mOTUpan (75% identity and 80% coverage thresholds) (Buck *et al.,* 2022) in genomes spanning the phylogenetic diversity of the wilt-pathogenic RSSC (n=23 genomes from the three species) as well as in “environmental” *Ralstonia* spp. from diverse habitats (n=29 genomes from 11 species; grey background on species tree). The presence and absence patterns of *in planta* fitness factors were mapped onto an approximately maximum likelihood species tree. Gene presence is indicated by rectangles colored according to functional classification, and gene absence is represented by white gaps. The species tree was generated using a KBase pipeline that constructs a tree with FastTree2 from a multiple sequence alignment of 49 conserved bacterial genes (Price *et al.,* 2010). Red asterisks indicate nodes with bootstrap support >0.8. Bold text highlights the RSSC species representatives used in the RB-TnSeq screens (Rpseu GMI1000, Rsol IBSBF1503, and Rsyz PSI07). The phylogenetic tree and presence/absence patterns were visualized using “ggtree” and “ggplot2” in R, and the figure was refined in Affinity Designer.

**Figure 3.**
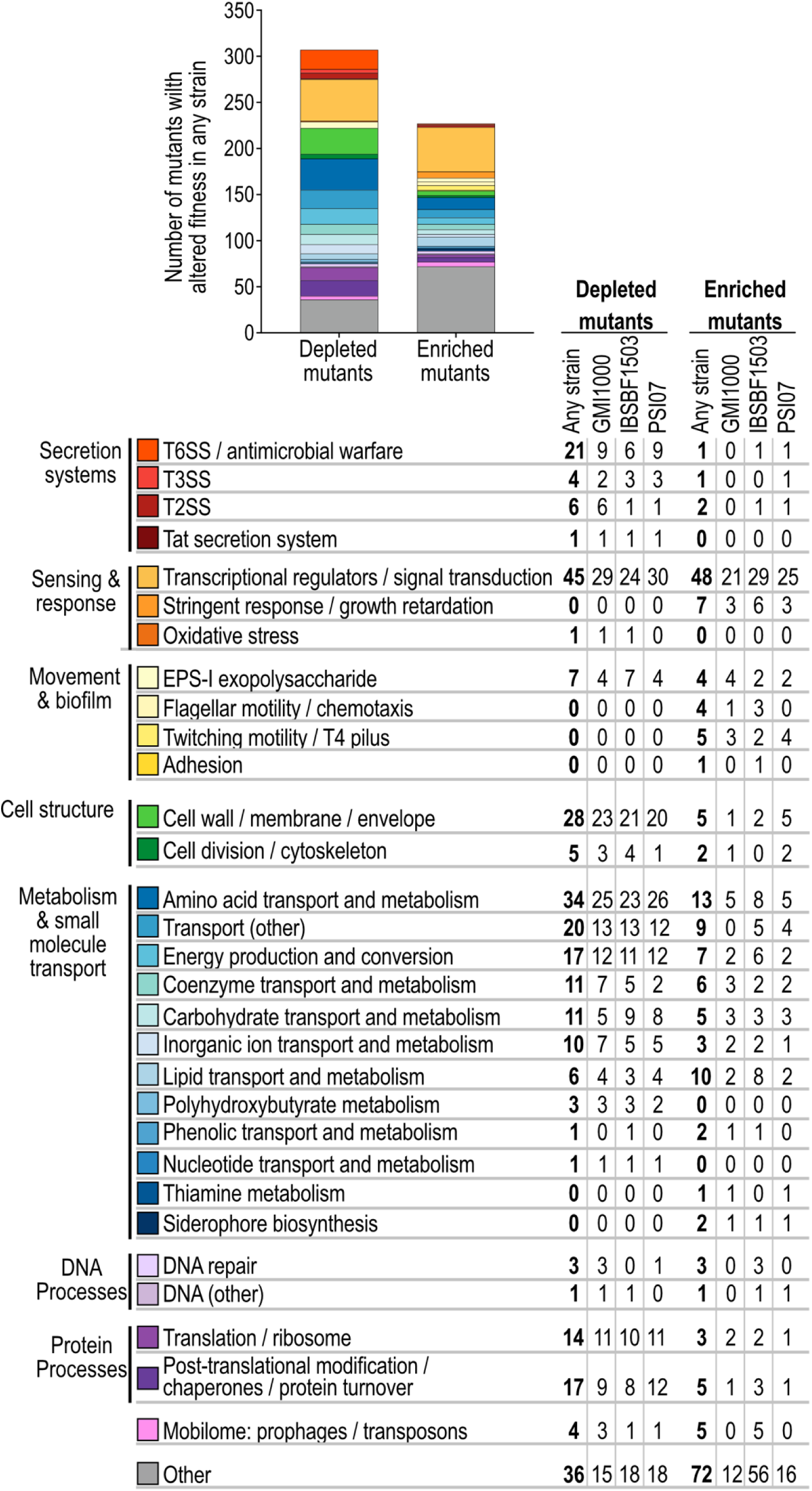
Functional classification of fitness-promoting and fitness-hindering genes affecting RSSC growth *in planta*. The fitness factors that affected *in planta* growth (|Fitness Score|<0.5) in Rpseu GMI1000, Rsol IBSBF1503, and Rsyz PSI07 were functionally classified. Categories were assigned using a combination of COG annotation (Tatusov *et al.,* 1997; Galperin *et al.,* 2025) and manual literature curation. Bar plots show the total number of mutants with depleted or enriched fitness in any strain with colors indicating functional categories.

Exploration of the resulting multi-species dataset revealed that orthologous genes often affected *in planta* fitness in the same direction across strains, although the effect sizes varied. For example, *phaR* mutant fitness scores ranged from -1.7 to -3.7 across the three strain backgrounds) (Table S4). However, some orthologous genes exhibited strain-specific fitness contributions, sometimes in opposing directions. For example, *dksA* mutant fitness score was -1.8 in Rspeu-GMI1000 and +3.3 in Rsol-IBSBF1503 strain. To quantify both the conservation and the natural variation of RSSC fitness factors, we used an intersection analysis (Upset plot) to visualize the overlap among strains and unique to one or two strains (Fig. 1d). The intersection analysis identified 88 genes that consistently promoted growth and 23 genes that consistently hindered growth. These numbers are likely an under-count of the shared fitness factors because individual RB-TnSeq libraries lacked barcoded mutants for orthologous gene families that promoted (n=29) or hindered (n=4) fitness in two of three strain backgrounds. For strain-specific genes, we found that 48, 44, or 47 genes uniquely promoted the growth of Rspeu-GMI1000, Rsol-IBSBF1503, or Rsyz-PSI07, respectively. Additionally, we found 27, 109, or 37 genes that uniquely hindered the growth of Rpseu-GMI1000, Rsol-IBSBF1503, or Rsyz-PSI07, respectively. Together, these data indicate both conserved and strain-specific genetic determinants underly RSSC pathogenic fitness.

### Many *in planta* fitness factors are unique to the RSSC wilt pathogens

To contextualize the genes that impact RSSC fitness *in planta*, we examined their conservation across the *Ralstonia* genus (Figs. 2, S9, and S10). We performed an evolutionary pangenome analysis using a phylogenetically balanced set of 23 RSSC genomes representing the three wilt-pathogenic RSSC species and 29 environmental strains from eleven other *Ralstonia* species isolated from diverse habitats (hereafter, “environmental *Ralstonia* spp.”). To focus comparisons on typical soil-borne RSSC vs. their relatives, we excluded the genome-reduced RSSC lineages that cause banana blood disease and Sumatra disease of clove [48].

Orthologs of the 307 growth-promoting and 227 growth-hindering genes were identified using 75% identity and 80% coverage thresholds with mOTUpan [44]. We organized the presence and absence of growth-promoting and growth-hindering genes by functional category and projected their distribution onto a phylogenetic tree of the strains (Fig. 2). Many factors that promoted (66.6%) *in planta* growth were well-conserved across the genus based on a threshold of being present in more than 95% of the 52 genomes. Using the same threshold, far fewer of the genes that hindered RSSC growth *in planta* were conserved genus-wide (35.1%). Across the functional categories, there were different proportions of genetic conservation (Figs. 2, S9, and S10). Several functional categories of fitness genes were universally or nearly universally conserved across the entire genus: protein processes (26/30 of growth-promoting and 8/8 of growth-hindering genes), DNA processes (3/4 of growth-promoting and 2/4 of growth-hindering genes), cell structure (26/33 of growth-promoting and 4/7 of growth-hindering genes). Within the metabolism and metabolite transport category, most growth-promoting genes were also highly conserved (97/114), whereas the growth-hindering genes had less conservation (32/58).

We were curious whether hierarchical clustering of gene presence/absence of fitness factor orthologs would recapitulate the phylogenetic relationships of the strains. Interestingly, hierarchical clustering of these orthologs largely recapitulated the phylogenetic relationships of the strains (Figs. S9 and S10). The hierarchical clustering cleanly separated the RSSC clade from the environmental clade. Moreover, the cladogram recapitulated the within-species relationships of RSSC strains accurately, except in the position of the RB-TnSeq strains. The RB-TnSeq strains had more singleton fitness factors, resulting in their placement as an outgroup to the RSSC branch.

Overall, these analyses suggest that while many fitness factors are broadly conserved across the genus, plant pathogenic RSSC may have also horizontally acquired a distinct repertoire of factors that support pathogenic success.

### Immunity genes involved in antimicrobial warfare are major, strain-specific determinants of *in planta* growth

Type VI secretion system (T6SS) immunity genes stood out as the strongest and most strain-specific fitness factors in our RB-TnSeq screens, suggesting that RSSC have evolved to actively engage in antimicrobial warfare while proliferating in their plant hosts (Fig. 4) [49]. To better understand the diversity of T6SS immunity genes, we recently compiled an atlas of T6SS genes the RSSC pangenome [50]. We classified dozens of auxiliary T6SS gene clusters (*aux1-aux25*) encoding a minimum of a VgrG spike protein, a toxin, and an immunity gene [50]. Here, we name the immunity genes as *rsi1* to *rsi25* for RSSC type six immunity gene from the *aux1* to *aux25* clusters.

**Figure 4.**
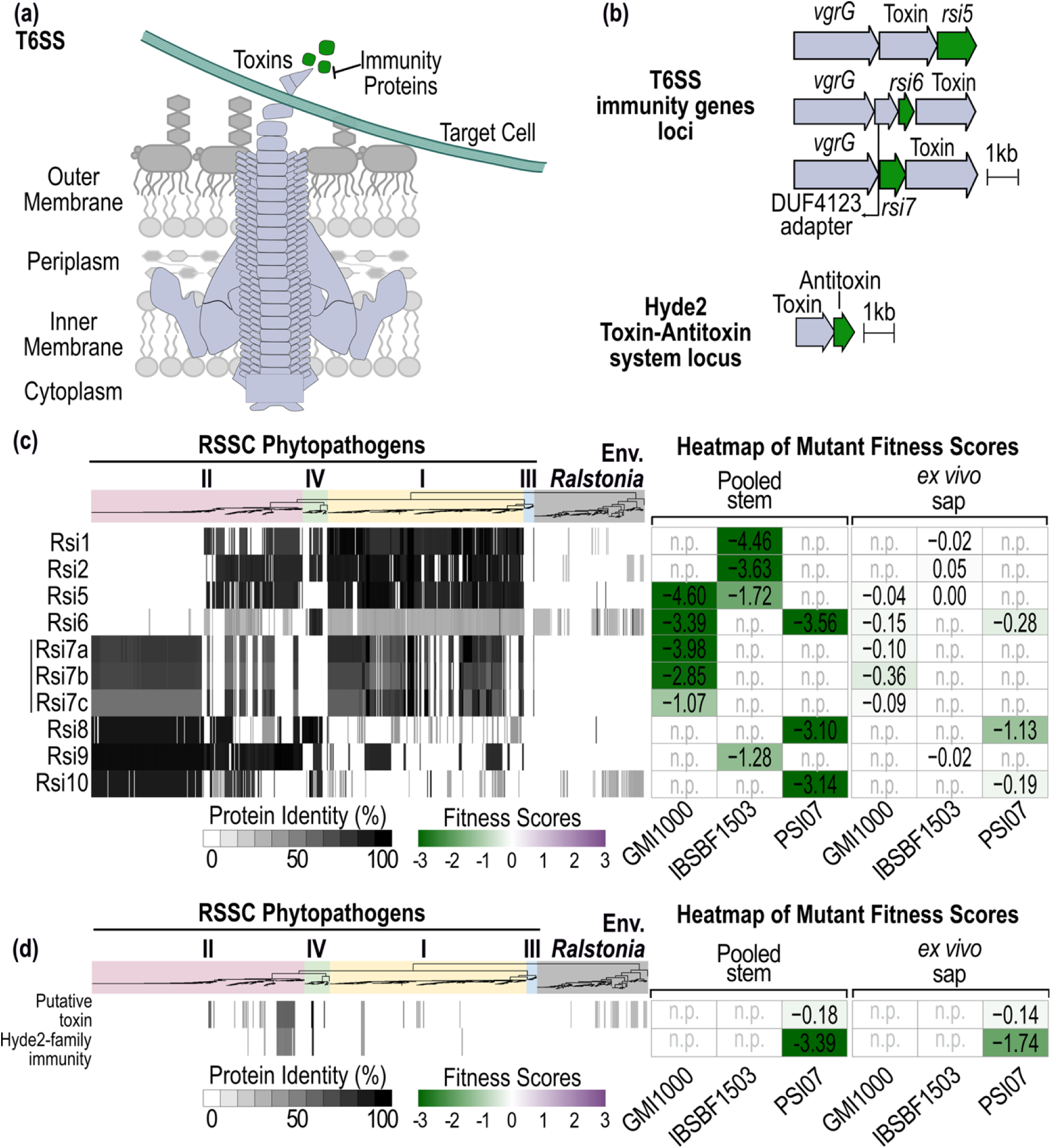
Type VI secretion system. (**T6SS) and HyDe2 immunity genes have strong and strain-specific promotion of RSSC growth *in planta*.** (**a**) Schematic representation of the T6SS delivering antimicrobial toxins into a kin cell, where cognate immunity proteins confer protection. (**b**) Genetic loci of three *vgrG-*linked immunity genes (in the Rpseu GMI1000 background) and a putative toxin/HyDe2 antitoxin system (in the Rsyz PSI07 background) (Levy *et al.,* 2017). (**c**) Phylogenetic distribution of canonical T6SS immunity proteins and (**d**) putative toxin/HyDe2 immunity proteins across the *Ralstonia* genus (621 plant-pathogenic RSSC genomes and 150 environmental *Ralstonia* spp. genomes). (**c-d Left**) The percent identity of the best BLASTp hit per genome with ≥60% coverage is shown as greyscale heatmaps below the phylogenetic trees. The phylogenetic tree was generated in KBase as previously described. Colored backgrounds denote phylotypes of plant pathogenic RSSC versus environmental *Ralstonia* strains. Phylogenetic trees and BLASTp results were visualized using iTOL (Letunic and Bork, 2021). (**c-d Right**) *In planta* and *ex vivo* xylem sap fitness score of homologs in RSSC species representatives (Rpseu GMI1000, Rsol IBSBF1503, and Rsyz PSI07) are shown as heatmaps ranging from green (mutants were depleted) to purple (mutants were enriched). “n.p.” indicates that a homolog was not present in the strain’s genome. and fitness. Heatmaps were generated using ‘pheatmap’ in R. The figure was refined in Affinity Designer.

In our *in planta* competitive fitness screens, each RSSC species representative required a different set of immunity genes for full growth in tomato stems. Rpseu-GMI1000 required *rsi5, rsi6,* and three *rsi7* paralogs (fitness scores: -1.1 to -4.6), Rsol-IBSBF1503 required *rsi1, rsi2*, *rsi5*, and *rsi9* (fitness: -1.3 to -4.6), and Rsyz-PSI07 required *rsi6*, *rsi8*, and *rsi10* (fitness: -3.1 to -3.6). The *rsi5* and *rsi6* homologs consistently promoted *in planta* fitness in both strains backgrounds where they occurred. In contrast, only one strain required *rsi1* and *rsi2* for growth *in planta* even though all three genomes contained *rsi1* and *rsi2* homologs (Fig. 4c). Except for *rsi8* in Rsyz-PSI07, the immunity genes were dispensable for RSSC growth in xylem sap (Fig. 4c).

We also identified a Hyde2-family protein in Rsyz-PSI07 that was required for normal *in planta* growth (Fig. 4d). This protein family was recently identified as being associated with toxin-antitoxin gene clusters, but its status as a toxin or immunity protein was not yet determined [51]. The Rsyz-PSI07 TnSeq results suggests the Hyde2 protein functions as an immunity protein.

Homologs of immunity genes were variably present across the RSSC (Fig. 4c and Table S4), consistent with our prior findings that these gene clusters have dynamic horizontal gene flow and that the immunity genes in Rpseu-GMI1000, Rsol-IBSBF1503, and Rsyz-PSI07 are non-orthologous xenologs [50]. Here, we show that homologs of these *in planta*-required immunity proteins were rare outside of the RSSC. Most *rsi* queries yielded few BLAST hits in the environmental *Ralstonia* genomes. When hits were present, most had low sequence identity (25-40%) to the RSSC queries, except for rare hits with over 60% identity. This pattern suggests that there is minimal-to-no gene flow of *aux* clusters between RSSC wilt pathogens and environmental *Ralstonia* species.

### Type III secretion system genes had complex impacts on fitness *in planta*

Multiple type III secretion system (T3SS) genes, predominantly transcriptional regulators or chaperones that aid effector secretion, impacted competitive *in planta* fitness of RSSC in positive or negative directions (Figs. 5, S11). The fitness impacts were specific to the *in planta* environment; the T3SS mutants grew like wildtype in xylem sap (Fig. S11).

**Figure 5.**
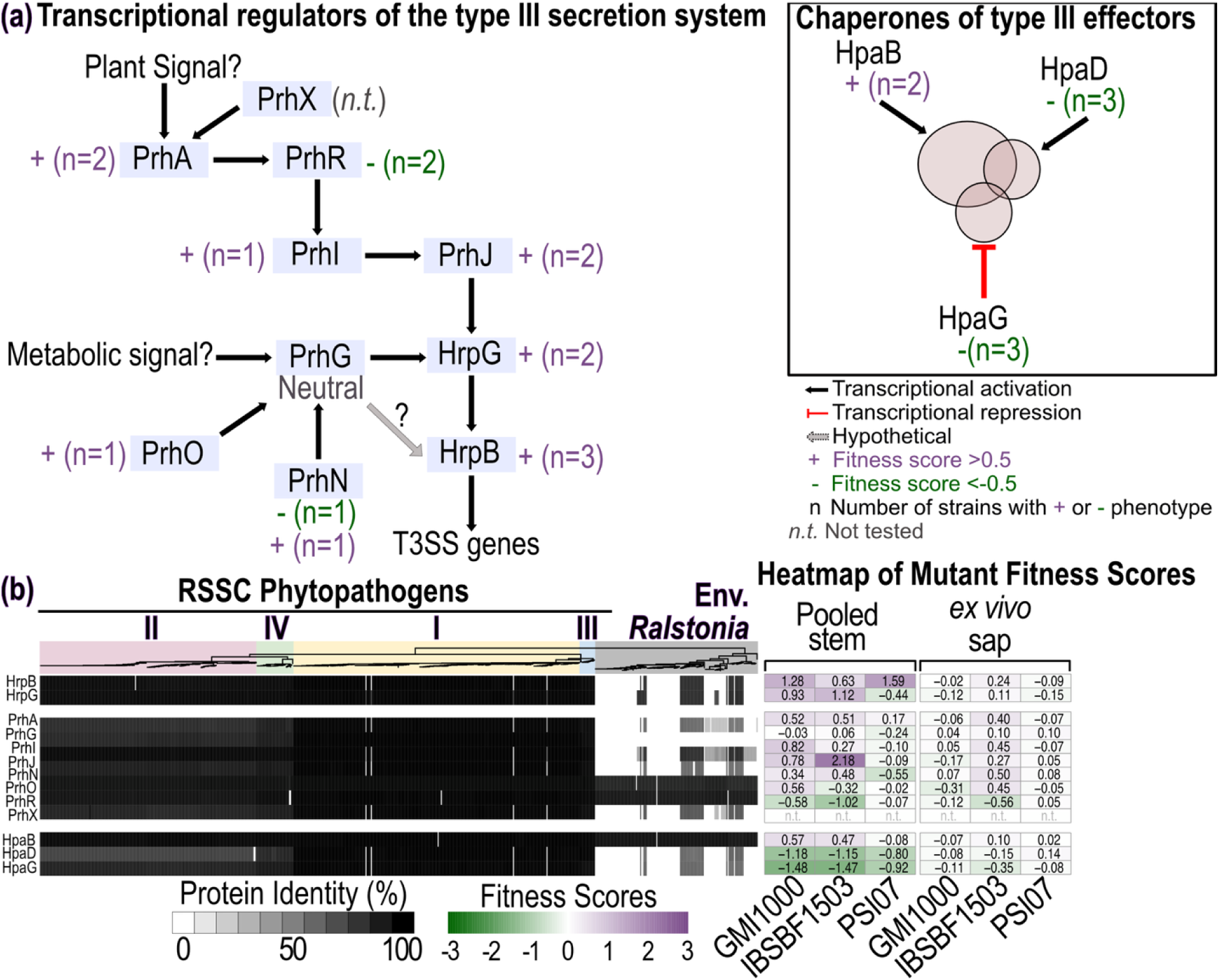
Transcriptional and post-translational regulators of the type III secretion system (T3SS) and effector secretion have complex *in planta* fitness patterns. (**a**) Schematic representation of the regulatory network controlling T3SS expression in Rpseu GMI1000 (adapted from Plener *et al.,* 2010). Plant-derived and metabolic signals are integrated through a hierarchical, branched regulatory network involving PrhA, PrhR/I/J, PrhG, and HrpG, ultimately activating the key regulator HrpB to regulate expression of T3SS genes. Black arrows indicate transcriptional activation, and grey arrows denote unresolved regulatory interactions. Chaperones of effector secretion, HpaB and HpaD, and the repressor HpaG are shown separately. Colored symbols next to each regulator indicate the mutants’ competitive fitness *in planta*: purple “+” denotes increased fitness (fitness score ≥ 0.5); green “-” denotes decreased fitness (fitness score ≤ -0.5). *“n”* indicates the number of strain backgrounds that showed the indicated phenotype(s). (**b Left**) Conservation of T3SS regulatory proteins across the *Ralstonia* genus (n=536 plant-pathogenic RSSC genomes and n=150 environmental *Ralstonia* spp. genomes). Proteins from Rpseu GMI1000 were queried, and the sequence identity of the best BLASTp hit with ≥60% coverage is shown as greyscale heatmaps below the phylogenetic tree. The phylogenetic tree was generated in KBase as previously described. Colored backgrounds denote phylotypes of plant pathogenic RSSC versus environmental *Ralstonia* strains. Due to a systematic annotation bias in which the Prokka implementation in KBase fails to annotate intact *prhA* genes in RSSC phylotype II genomes, so 85 phylotype II genomes were excluded; the retained 202 phylotype II genomes are RefSeq assemblies annotated using NCBI PGAP. The phylogenetic tree and BLASTp results were visualized using iTOL (Letunic and Bork, 2021). (**b Right**) *In planta* and xylem sap fitness score of orthologs in RSSC species representatives (Rpseu GMI1000, Rsol IBSBF1503, and Rsyz PSI07) are shown as green-to-purple heatmaps on the right. “n.t.” indicates not tested due to absence of corresponding mutants in the RB-TnSeq library. Fitness heatmaps were generated using ‘pheatmap’ in R. The figure was refined in Affinity Designer.

Expression of RSSC T3SS genes is activated by a well-characterized, transcriptional regulatory network where three branches converge to activate HrpG (Fig 5a) [52–56]. HrpG activates expression of *hrpB*, and HrpB specifically binds the *hrpII* boxes in promoters of T3SS operons [57].

HrpB had a consistent growth-hindering phenotype across the three species representatives (fitness: +0.6 to +1.6), but multi-strain fitness patterns for other T3SS transcriptional regulators were more complex with fitness scores in the Rsyz-PSI07 background often differing from shared patterns in the other two strains (Fig. 5b). Most of the upstream T3SS expression activators also had growth-hindering phenotypes in the Rpseu-GMI1000 and Rsol-IBSBF1503 backgrounds: HrpG (fitness: +0.9 and +1.1), PrhA (fitness: +0.5 and +0.5), PrhJ (fitness: +0.8 and +2.2), and a trend in PrhN (fitness: +0.3 and +0.5). PrhI (fitness: +0.8) and PrhO (fitness: +0.6) also hindered growth specifically in the Rpseu-GMI1000 background. PrhR had an opposing phenotype (fitness: -0.6 in Rpseu-GMI1000 and -1.0 in Rsol-IBSBF1503 backgrounds) (Fig. 5b), even though it is also an activator of T3SS expression. Of these regulators, HrpG and PrhN conferred Rsyz-PSI07-specific, growth-promotion phenotypes (HrpG fitness: -0.4; PrhN fitness: -0.6), while the others had neutral impacts to fitness.

Two chaperones that deliver overlapping sets of effectors to the T3SS injectosome, HpaG and HpaD [14,58], consistently promoted *in planta* growth with mutants in all strain backgrounds having fitness scores of -0.8 to -1.5. In contrast, HpaB, the third T3 effector chaperone [58], imposed minor growth hindrance in two strains (fitness: +0.5 in Rsol-IBSBF1503 and +0.6 in Rsyz-PSI07). Although these chaperones control secretion of different repertoires of effectors, none of the effector mutants had altered fitness in these RB-TnSeq screens. Multiple structural proteins comprising the core export apparatus (HrpD, HrcN, HrpF, HrpH, and HrpK) mildly promoted fitness of strain Rsyz-PSI07 (fitness: -0.5 to -0.9) (Fig. S11c).

Although the T3SS is a major virulence factor that is known to be in the core RSSC genome, the phylogenetic conservation of this pathogenicity factor across the *Ralstonia* genus has not been explored to our knowledge. Most of the RSSC T3SS structural genes and regulators are encoded in a >35 kb cluster on the megaplasmid (Fig. S11a). BLASTp searches identified homologs of these clustered T3SS genes in almost all RSSC genomes. The T3SS was not present in several species: *R. thomassii*, *R. pickettii sensu stricto*, *R. insidiosa*, or *R. flaminis* genomes.

However, high identity T3SS orthologs were identified in 27% of environmental *Ralstonia* genomes: all *R. mojiangensis* (n=6 of 6), *R. holmesii* (n=5 of 5), *R. chuxiongensis* (n=4 of 4), *R. wenshanensis* (n=6 of 6), and *R. edaphis* (n=3 of 3) as well as some *R. mannitolilytica* (n=13 of 26) and *R. flatus* (n=1 of 3). In these environmental *Ralstonia* genomes, the T3SS blast hits were present in a gene cluster that was syntenic with the RSSC cluster. Overall, phylogenomics showed that multiple species of environmental *Ralstonia* encode T3SS genes with high identity to RSSC queries (Fig. S11a).

Four of the regulators of T3SS gene expression are encoded outside of the RSSC T3SS gene cluster, and these fitness factors exhibited distinct conservation patterns. The three encoded on the chromosome (*prhN, prhO,* and *prhX*) were universally conserved across the *Ralstonia* genus (Figs. 5b, 11c). In contrast, the megaplasmid-encoded *prhG* was specific to RSSC and absent from environmental *Ralstonia* genomes regardless of their T3SS status.

### The type II secretion system (T2SS) components but not T2SS cargo contributed to fitness

*Ralstonia* uses a type II secretion system to secrete environment-remodeling effector proteins into the extracellular space (Fig. S12), and T2SS mutants are known to have a virulence defect [20,59]. RB-TnSeq revealed that T2SS promoted *in planta* growth, with most consistent phenotypes for Rpseu-GMI1000 mutants (Fig. S12c). Strain Rsol-IBSBF1503 and Rsyz-PSI07 each had one T2SS gene associated with growth promotion (Fig. S12c). The T2SS gene cluster was conserved across the genus with syntenic gene organization and high sequence identity of orthologous genes (Fig. S12b-c). The known cargo of the RSSC T2SS had neutral impacts on fitness: Egl and CbhA cellulases, pectinases (PehA, PehB, PehC, and Pme), and NucA and NucB DNases [20,60,61].

### EPS-I biosynthesis genes have complex fitness contributions

Mutants involved in production of the biophysically unique EPS-I exopolysaccharide virulence factor had complex patterns of fitness effects (Fig. S13a) [32,40]. Mutants lacking the XpsR positive regulator of *eps* gene expression (RS_RS21960) [62] were enriched specifically *in planta* (Stem fitness: +0.9 to +1.9; xylem sap fitness: +0.1 to +0.2) (Fig. S13b). In all RSSC backgrounds, strong *in planta* fitness defects were exhibited by mutants lacking the EpsE inner membrane export protein (fitness: -1 to -2.2), the RS_RS21980 O-acetyltransferase (fitness: -1.7 to -3.1), or the RS_RS21975 aminotransferase (fitness: -1.2 to -3.1); these mutants had milder fitness defects in xylem sap (fitness: -0.2 to -1.0). Additionally, mutants lacking the EpsF inner membrane export protein, the GT4-family glycosyltransferase (RS_RS22010), the second O-acetyl transferase (RS_RS22005), the dehydrogenase (RS_RS22000), and the heparinase-family protein (RS_RS21995) had mild fitness defects of similar magnitudes *in planta* and in xylem sap (−0.1 to -1.1) (Fig. S13b). Conversely, Rpseu-GMI1000 mutants lacking the two epimerases (EpsC and RS_RS21965), the EpsD UDP-N-acetyl-aminosugar dehydrogenase (RS_RS22025), and the WcaJ-like glycosyltransferase (RS_RS21985) were mildly enriched *in planta* (fitness: +0.1 to +0.7) and had wildtype-level fitness in xylem sap. These contrasting fitness patterns are peculiar. It is possible that certain mutations relieve the metabolic burden of EPS-I production [63], whereas other mutations cause deleterious disruption to metabolic flux, leading to fitness defects.

### RSSC must maintain an effective cellular barrier *in planta*

Gene products that preserve integrity of the bacterial envelope promoted RSSC growth *in planta* (Fig. 6a-c). In most strain backgrounds, mutations in lipopolysaccharide modification genes (rfbA, rfbF, *rfaD*, *lgtF*, RS_RS03460, RS_RS03465, and RS_RS11045) strongly reduced fitness both *in planta* and in xylem sap down to fitness scores of -4.3 (Fig. 6d), consistent with previous work [64]. Among these seven genes, only *rfaD* was a core gene across the *Ralstonia* genus; for the other LPS fitness factors, the strong conservation in the RSSC clade and absence in the environmental *Ralstonia* clade suggests that these gene products are adaptations to the wilt-pathogenic lifestyle of the RSSC (Fig. 6d).

**Figure 6.**
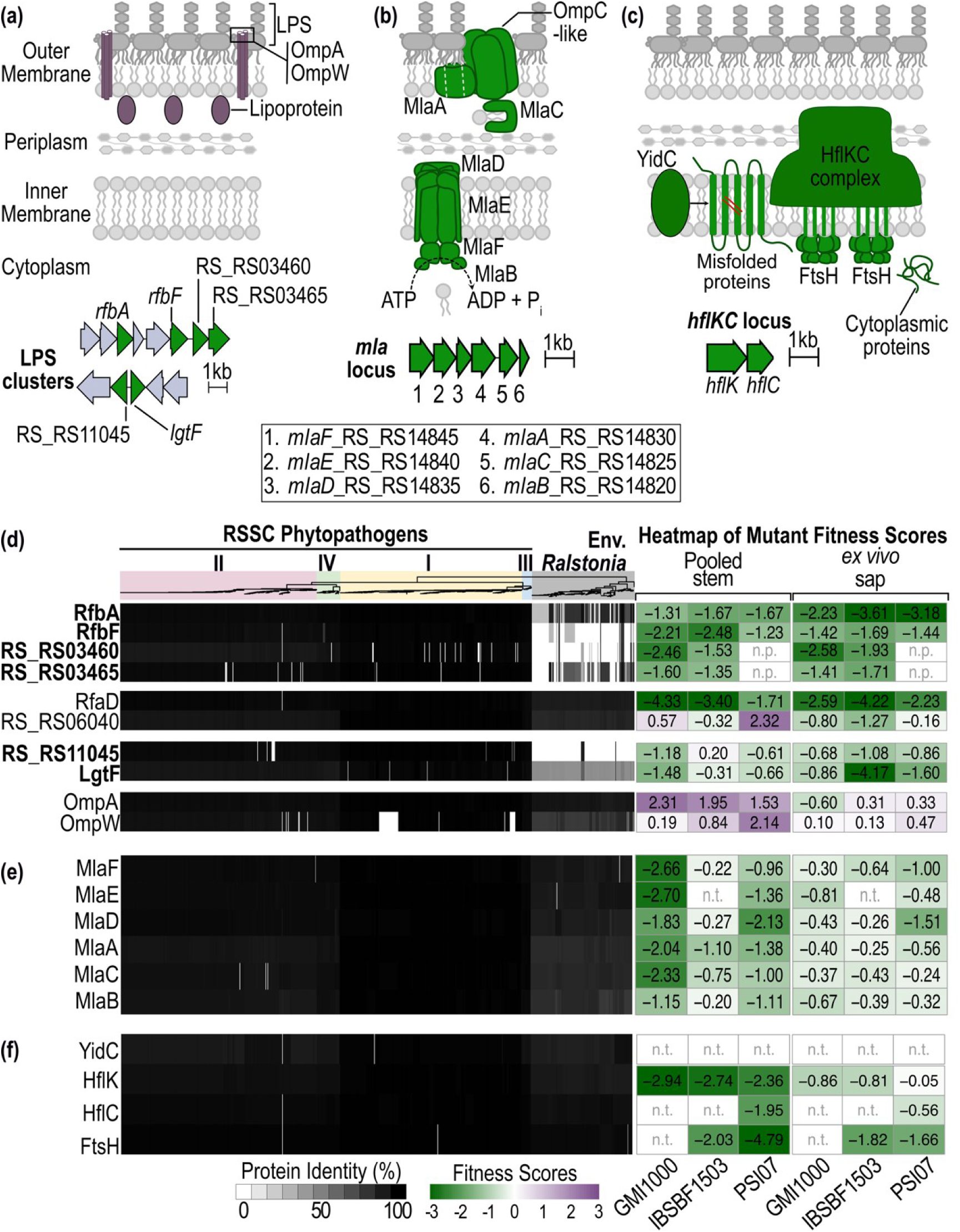
Genes involved in envelop integrity are consistently required for *in planta* growth of wilt-pathogenic RSSC. (**a-c**) Schematic representations of envelop traits that affect *in planta* fitness. (**d**-**f Left**) Conservation of outer membrane-related and LPS biosynthesis proteins, the maintenance of lipid asymmetry (Mla) system, and the FtsH-HflKC misfolded protein degradation system across the *Ralstonia* genus. The phylogenetic tree of 621 plant-pathogenic RSSC strains and 150 environmental *Ralstonia* spp. strains was generated in KBase as previously described. Colored backgrounds denote phylotypes of plant pathogenic RSSC versus environmental *Ralstonia* strains. Percent identity of the best BLASTp hit per genome with >60% coverage is shown as greyscale heatmaps below the phylogenetic tree. Phylogenetic trees and BLASTp results were visualized using iTOL (Letunic and Bork, 2021). (**d-f Right**) *In planta* and xylem sap fitness score of orthologs in RSSC species representatives (Rpseu GMI1000, Rsol IBSBF1503, and Rsyz PSI07) are shown as green-to-purple heatmaps on the right. “n.p.” indicates not present due to missing ortholog. “n.t.” indicates not tested due to absence of the corresponding mutant in the RB-TnSeq library. Fitness heatmaps were generated using ‘pheatmap’ in R. The Figure was refined in Affinity Designer.

RSSC required the maintenance of outer membrane lipid asymmetry (Mla) system for full *in planta* growth, with *mlaFEDACB* mutants in the Rpseu-GMI1000 background exhibiting the strongest defects (fitness: -1.1 to -2.7) (Fig. 6b,e). To maintain an effective barrier, phospholipids that are mislocalized on the outer leaflet of the outer membrane are extracted by MlaA, transferred to the periplasmic carrier MlaC, and delivered to the inner-membrane ABC transporter MlaFEDB for reinsertion into the inner membrane. The *mla* mutants had more pronounced fitness defects when growing *in planta* than their mild fitness defects when growing in xylem sap. Phylogenetically, the Mla system was conserved across the *Ralstonia* genus with higher protein sequence conservation in MlaFED (≥85%) than MlaACB (≥77%) (Fig. 6e).

The HflKC-FtsH quality control system regulates proteolysis of misfolded inner-membrane proteins as well as multiple envelope-associated regulatory factors. This proteolytic system was more strongly required for RSSC growth *in planta* than in xylem sap across all strain backgrounds, although *hflC* and *ftsH* mutants were not represented in all mutant libraries (Fig. 6c,f). The HflKC-FtsH complex is phylogenetically conserved across the genus with high sequence identity (≥89.6%).

In contrast, several envelope-localized proteins conferred fitness costs *in planta*. Although an OmpA-family porin (RS_RS11950) was universally conserved in the genus, mutants lacking this gene had a growth advantage *in planta* (fitness: +1.5 to +2.3) despite neutral fitness in *ex vivo* xylem sap (fitness: -0.6 to +0.3), suggesting that this porin may allow harmful import of infection-induced phytodefense chemicals (phytoalexins) into the periplasm. A putative periplasmic peptidoglycan hydrolase (RS_RS06040) had contrasting fitness impacts *in planta* (increased fitness) versus in xylem sap (decreased fitness) in two strain backgrounds (Fig. 6d).

### The stringent response may curb RSSC growth *in planta*

Bacteria sense nutrient limitation and prepare for stationary phase through multiple pathways, including the stringent response [65]. Multiple stringent response and stationary phase genes hindered *in planta* growth of one or more RSSC strains: *relA, spoT, dksA, rsfS, rpoS,* and two HipA-family toxins (RS_RS12575 and RS_RS12575) (Table S4). Mutants in *rpoS, spoT, relA,* and RS_RS12575 had enriched *in planta* abundance across strain backgrounds; the *rpoS* and RS_RS12575 mutants showed larger magnitude phenotypes in Rpseu-GMI1000 and Rsyz-PSI07 vs. Rsol-IBSBF1503. Phenotypes for the other mutants (*rsfS, dksA,* and RS_RS07280) were strain-dependent with *in planta* phenotypes in opposing directions for some mutants (Tables S4 and S5). Notably, *dksA* mutants had a defect *in planta* in one strain background (Rpseu-GMI1000 fitness: -1.8), wildtype-level *in planta* growth in the Rsyz-PSI07 background but strongly enhanced competitive *in planta* growth in the Rsol-IBSBF1503 background (fitness: +3.3). Most of the stringent response mutants largely had wildtype-level growth in xylem sap, except for *dksA* mutants (xylem sap fitness: -0.8 to -1.2) [23].

Most stringent response genes were phylogenetically conserved across the *Ralstonia* genus (*relA*, *rsfS*, *rpoS*, *spoT, dksA,* and RS_RS12575), while the weaker phenotype HipA family-encoding gene (RS_RS07280) was core to the RSSC and rare in environmental *Ralstonia* genomes.

### Metabolic functions like sugar utilization, polyhydroxybutyrate metabolism, and phosphate uptake contributed to competitive fitness *in planta*

Metabolism and small molecule transport represented the largest functional group of genes that promoted growth (114/307 genes) and hindered growth *in planta* (58/227) (Figs. 1-3). As expected, most metabolic functions contributed similarly to RSSC fitness *in planta* and in xylem sap, such as the growth defects exhibited by amino acid biosynthesis mutants [23] (Table S4).

Sugar utilization pathways supported RSSC carbon acquisition *in planta.* Although sugars are not the main carbon sources for the RSSC *in planta* [17,66,67], RB-TnSeq provides additional evidence that sucrose catabolism via the *scrRAB* genes contributes to pathogen fitness [11,68]. Specifically, *scrB* mutants had defects across all RSSC strains (fitness: -0.9 to -3.7), and all three *scr* operon genes were required for *in planta* fitness in the Rsol-IBSBF1503 background. The *scrRAB* gene cluster was broadly conserved across the RSSC while absent from environmental *Ralstonia* genomes (Fig. S9, and Table S5), suggesting sucrose utilization is a pathogenic metabolic strategy. Glucose catabolism genes were also important *in planta:* glucose-6-phosphate isomerase *(pgi* fitness: *-*2.2 to -3.2), glucose-6-phosphate dehydrogenase *(zwf* fitness: -0.3 to -3.2) and the Entner-Doudoroff pathway gene *edd* (fitness: -0.1 to −3.3) all promoted in *planta* growth in multiple strains, with milder xylem sap defects. Additionally, an RSSC-specific EamA/RhaT family carbohydrate transporter (RS_RS20405) consistently promoted the pathogens’ growth exclusively *in planta* (Fig. S9, Tables S4, and S5).

Synthesis of polyhydroxybutyrate (PHB), an energy and carbon storage molecule, was important for *in planta* fitness. PHB enzymes and regulators had stronger fitness effects *in planta* (*phaR* average fitness: -2.6; *phbB*: -0.8; and *phbC*: -2.1) than in xylem sap (average fitness: -0.5, -0.5, and -1.7, respectively) (Fig. S9, Tables S4, and S5). The *phcCABR* gene cluster is phylogenetically conserved across the genus.

Phosphate uptake, via the PstSCAB transporter, contributed to RSSC growth *in planta* (Fig. S14). *pst* mutants in all strain backgrounds had reduced fitness *in planta* (fitness: -0.5 to - 2.5), while PhoU had increased fitness *in planta* (fitness: +1.3). Neither the Pst system nor PhoU affected RSSC growth in xylem sap. The *pstSCAB-phoUBR* gene cluster is phylogenetically conserved across the genus (Fig S14b).

### Signal transduction and transcriptional regulators impact RSSC competitive fitness *in planta*

Sensing, signal transduction, and transcriptional regulators constituted a major fraction of genes affecting *in planta* fitness: 46/307 of growth-promoting and 55/227 of growth-hindering factors (Figs. 1-3). Many regulators impacted *in planta* growth in at least one RSSC strain background (Figs. 1, 3, and Table S4), including the aforementioned *hrpB*, *hrpG, prhA, prhI, prhJ, prhN, prhO, scrR, phoU, xpsR, phaR,* and the stringent response regulators.

Among the characterized global virulence regulators, several impacted RSSC fitness in expected ways while others had unexpected phenotypic effects. The global regulator EfpR, previously discovered via serial *in planta* passage evolution experiments with Rpseu-GMI1000 [12,18,69], conferred increased *in planta* fitness in only Rpseu-GMI1000 and Rsol-IBSBF1503 (fitness: +5.3 and +1.5, respectively) the paralogous EfpH conferred increased fitness in all three strain backgrounds (fitness: +0.5 to +2.4). Neither EfpR nor EfpH impacted fitness in sap [23].

EfpR/H are pathogen-specific genes although environmental *Ralstonia* genomes encoded rare, *efp-*like genes that lacked orthologous neighborhoods (Table S5 and data not shown). Surprisingly, mutations in the VsrAD and VsrBC two component system regulators [70] caused unexpected cheating phenotypes where *vsrAD* mutants had strongly increased fitness (+6.0 to +7) and *vsrBC* mutants had moderately increased fitness (+0.6 to +1.7). Both VsrAD and VsrBC are conserved across the entire genus (Table S5). Quorum sensing mutants had strain-dependent phenotypes *in planta*. In addition to what has been shown for *in planta* fitness of *phcA* mutants in the Rpseu-GMI1000 background [46] (fitness: +2.1), Rsol-IBSBF1503 *phcA* mutants also cheated *in planta* (fitness: +2.7), but Rsyz-PSI07 mutants had an contrasting loss-of-fitness phenotype *in planta* (fitness: -1.8); these patterns were also seen in *ex vivo* xylem sap [23]. In all three backgrounds *phcR* mutants had reduced fitness *in planta* (fitness: -0.6 to -2.5) that was not exhibited in xylem sap [23]. *phcS* mutants had reduced fitness in Rpseu-GMI100 (fitness: -1.1) and Rsyz-PSI07 (fitness: -2.2) backgrounds, despite increased fitness in the Rsol-IBSBF1503 background (fitness: +0.6); *phcS* mutants did not show fitness defects in xylem sap [23]. These quorum sensing genes are universal in the RSSC and widely prevalent, but non-universal, in the environmental genomes (72-86%) (Table S5).

RpoN1 [71,72] and NarL, two nitrogen metabolism regulators, had opposite contributions to *in planta* growth with presence of *rpoN1* promoting growth of all strains (fitness: -0.8 to -1.4) and presence of *narL* hindering growth of Rsol-IBSBF1503 (fitness: +0.7). RpoN1 is core to the genus while *narL* is enriched in the RSSC; all RSSC encode *narL* while only 10.3% of environmental *Ralstonia* genomes encode an ortholog (Fig. S9 and Table S5).

A putative oxidative-stress responsive two-component system (PrrAB) specifically promoted *in planta* growth across all strain backgrounds (fitness: -1.2 to -2.7). This unstudied two-component system is conserved across the *Ralstonia* genus (Table S5).

Of the unnamed regulators, five had strain-independent impacts on competitive growth *in planta,* despite relatively neutral impacts in xylem sap: RS_RS24715 (fitness: +1.5 to +3.3), RS_RS25105 (fitness: +1.8 to +2.5), RS_RS25110 (fitness: +1.6 to +2.4), RS_RS18630 (fitness: +1.1 to +1.9), RS_RS16070 (−0.8 to -1.3). Of these, three were conserved across the *Ralstonia* genus (RS_RS25105, RS_RS25110, RS_RS18630). The RS_RS16070 TetR-family regulator was sporadic across the genus, present in 69.6% of RSSC and 34.5% of environmental *Ralstonia*. The pathogen-specific RS_RS24715 helix-turn-helix transcriptional regulator was only present in RSSC genomes.

### Several hypothetical proteins of unknown function impact RSSC fitness

We identified an interesting set of fitness factors lacking known annotations that contribute to RSSC fitness during *in planta* growth. Across all strain backgrounds, these non-annotated fitness factors consistently promoted (RS_RS02305, RS_RS10075, RS_RS10430, RS_RS10435, and RS_RS12580) or consistently hindered (RS_RS01160) RSSC growth *in planta.* Three of these uncharacterized genes displayed strong *in planta* fitness effects while having weak, neutral, or variable effects in xylem sap, including RS_RS12580 (fitness: -0.6 to -1.6 *in planta* versus 0 to +4 in xylem sap), RS_RS01160 (fitness: +2.1 to +3 *in planta* versus +0.2 to +0.5 in xylem sap), and RS_RS08165 (fitness: +0.5 to +1.7 *in planta* versus +0.5 to -0.2 in xylem sap). Other fitness factors of unknown function were encoded in only one RB-TnSeq strain; seven factors promoted the strain’s growth (RS_RS25400, RS_RS25615, RALBFv3_RS20925, RALBFv3_RS24385, RPSI07_RS20725, RPSI07_RS24665, and RPSI07_RS03950), and 13 hindered the strain’s growth (RS_RS25830, RALBFv3_RS08590, RALBFv3_RS16780, RALBFv3_RS19305, RALBFv3_RS19685, RALBFv3_RS20085, RALBFv3_RS20920, RALBFv3_RS22620, RALBFv3_RS23095, RALBFv3_RS23680, RALBFv3_RS24045, RPSI07_RS24195, and RPSI07_RS24990). Genes with these patterns of phylogenetic distribution and fitness contributions may contribute to phenotypic heterogeneity of the RSSC.

## Discussion

Understanding how RSSC pathogens achieve fitness within the plant xylem is central to explaining the rapid progression of bacterial wilt diseases. By integrating evolutionary genomics with quantitative, multi-strain forward genetic screens performed directly *in planta*, this study reveals that RSSC pathogenic and ecological success emerges from the interplay of conserved physiological resilience, lineage-specific adaptations, and being pre-emptively prepared for antimicrobial warfare during plant infection. Comparing *in planta* fitness landscapes with growth in *ex vivo* xylem sap showed that many physiological traits required for growth in sap remain important *in planta*, while additional genetic requirements emerge under host-specific pressures. Our results are consistent with *Xanthomonas* and *Pseudomonas syringae* studies that compared multiple infection-relevant conditions and found substantial overlap in fitness requirements, while consistently revealing that the *in planta* environment demands a more expansive repertoire of fitness determinants [73,74]. Xylem sap represents a reductionist plant-like environment containing constitutive chemical defenses such antimicrobial phenolics and defense-related proteins [75]. In contrast, the *in planta* environment exposes mutant libraries to induced defenses deployed following immune perception [75]. The quantitative fitness data demonstrates that even in these susceptible tomato plants, RSSC pathogens require a plethora of sensory and regulatory proteins and envelope maintenance systems to be resilient against induced host defenses.

We initially hypothesized that the pathogens would require identical metabolic programs for growth in tomato stems that they required for growth in the *ex vivo* xylem sap. For most metabolic pathways, the fitness results were consistent with this null hypothesis [23]. However, mutants lacking certain metabolic genes showed stronger or exclusive fitness defects *in planta*: phosphate uptake, polyhydroxybutyrate metabolism, sucrose utilization, and glycolysis. These results indicate one of two phenomena: (1) availability of phosphate and sugars in xylem sap changes during infection (supported by [68]) and/or (2) RSSC pathogens have less access to phosphate and higher access to sugars (or lower access to their preferred carbon sources, amino acids) after the bacterial pathogens ooze into the stem apoplast. Future investigation into the spatial landscape of nutrient availability and metabolic programs could shed light on whether these pathogens display phenotypic heterogeneity of their metabolic programs *in planta*.

To explore how the RSSC evolved to be aggressive wilt pathogens, we investigated the phylogenetic distribution of fitness factors across the *Ralstonia* genus. These evolutionary comparisons allowed us to classify traits as putatively ancestral if the genes were conserved across the genus, putatively linked to emergence of wilt pathogenesis if conserved across the RSSC while absent from genomes of the environmental *Ralstonia* spp., or lineage-specific adaptations if present sporadically in the RSSC. Our data are consistent with prior inferences that the Phc quorum sensing system is sporadically present in genomes of the pathogens’ evolutionary neighbors [76], and that EPS-I biosynthesis may have been a key evolutionary event for wilt pathogenesis [32].

The phylogenetic distribution pattern of the T3SS was unexpected—a high-identity, syntenic T3SS gene cluster was present in multiple lineages of environmental *Ralstonia* that do not cause wilt disease. Future studies should investigate the evolutionary history of the RSSC T3SS, a monumentally important virulence factor that allows RSSC pathogens to manipulate living plant cells. Was the T3SS an ancestral trait to the *Ralstonia* genus that was subsequently lost? Alternatively, the phylogenetic distribution could be consistent with the RSSC ancestor acquiring the T3SS after it diverged from the other *Ralstonia* species with subsequent horizontal transmission of the T3SS gene cluster to ecologically sympatric *Ralstonia* species. The multi-strain fitness patterns of T3SS regulators did not neatly align with the regulatory model developed from targeted mutagenesis approaches in Rpseu-GMI1000 [77,78]. Since the regulators converge on regulating *hrpG* expression and HrpG regulates multiple functions other than T3SS [10], it is possible that the fitness signatures of these mutants are reflecting their role in regulating other cell functions.

Among the lineage-specific fitness determinants identified, T6SS immunity genes and the Hyde2 antitoxin emerged as major requirements for RSSC strains’ full growth *in planta.* These immunity proteins were variably distributed across the RSSC phylogeny, consistent with diversification of toxin-immunity repertoires across the RSSC complex, as we previously showed with a curated set of well-assembled genomes [50]. The immunity mutants’ fitness defects suggest that RSSC deploy their T6SS-secreted toxins *in planta*, leading to the immunity mutants experiencing toxin-mediated self-harm. Most immunity mutants had no fitness defect in *ex vivo* xylem sap or rich broth or minimal medium broth [23]. The specific *in planta* requirement of these immunity proteins suggests that RSSC have evolved to wield the cognate T6SS toxins specifically in the *in planta* environment. Due to the widespread usage of the T6SS as a weapon that targets competitor bacteria [49], we hypothesize that the main target of the RSSC T6SS are competing bacteria in the phytobiome. Because the *in planta* TnSeq screens involved inoculation of a single RSSC strain background, and the xylem of young, growth chamber-grown tomato plants likely have few-to-no-endophytic microbes present, we infer that RSSC deploy their T6SS toxins *in planta* even in the absence of known competitors. However, existing data from this study or the literature cannot rule out the role of T6SS toxins as causing lytic damage to host cells because some bacteria target eukaryotic cells with their T6SS [79]. A better understanding of the RSSC T6SS is warranted.

Traditional approaches to understanding pathogenesis have attempted to strictly divide pathogen traits into “virulence factors” vs. “factors that generally contribute to microbial growth”, whereas the modern paradigm is that pathogen virulence is integrated with pathogen physiology. Thus, although we identify certain traits as putatively ancestral to pathogenesis, the quantitative fitness data demonstrates that many ancestral traits may be essential pre-requisites for pathogenesis to have emerged. For example, the T2SS secretion system appears to be an ancestral trait even though the secretion system is essential for the pathogen to remodel the *in planta* physical environment with extracellular cellulases, pectinases, and nucleases [20,21,60,80]. Long-term ecological success of RSSC pathogens likely benefits from ancestral physiological traits. Since RSSC pathogens have a feast-vs-famine life cycle where they grow prolifically *in planta* in between periods of prolonged dormancy in the soil, we infer that the reduced competitive fitness of Stringent Response mutants is a short-term trade-off that promotes their ability to prepare for survival after killing their plant hosts.

Intriguingly, the number of mutants with increased competitive fitness varied considerably by RSSC strain background. There were 109 genes that hindered Rsol-IBSBF1503 growth *in planta*, while only 37 and 27 genes that hindered *in planta* growth of Rsyz-PSI07 and Rpseu-GMI1000, respectively. One potential explanation may lie in the host-of-isolation for these strains— Rpseu-GMI1000 and Rsyz-PSI07 were both isolated from wilting tomato plants (Solanaceae family), presumably during an outbreak of bacterial wilt disease in a monoculture field. In contrast, Rsol-IBSBF1503 was isolated in Brazil from wilting cucumber plants (Cucurbitaceae family) [81]. It is possible that the Rsol-IBSBF1503 strains’ ancestors had not infected tomato plants, so there were many routes to becoming more genetically adapted to the tomato stem environment. Taking advantage of the Ralstonia Wilt Dashboard [82], we investigated the reported hosts-of-isolation for the phylotype IIB sequevar 4 (IIB-4) lineage that includes Rsol-IBSBF1503 [81]. Although tomato was the fourth most reported host -of-isolation for IIB-4 strains, no IIB-4 strains are present among the 168 reported Brazilian tomato isolates reported from 13 studies [83–95]. Thus, it is possible that Rsol-IBSBF1503 is highly virulent despite not being specifically adapted to tomato.

Any high-throughput dataset is susceptible to false positives and false negatives. To evaluate potential error in our dataset, we compared our replicated *in planta* RB-TnSeq results with a prior study profiled a single biological replicate of a Rpseu-GMI1000 TnSeq library in a different susceptible tomato cultivar (Figs. S15 and S16) [18]. We observed concordant effects for 59 genes in Rpseu-GMI1000 (58 genes whose disruption reduced *in planta* fitness and one gene whose disruption increased fitness). Using the replicate variation in our dataset as a benchmark, we infer that the prior single-replicate screen likely missed a subset of true effects (potential false negatives: 149 genes promoting growth and 68 genes hindering growth in Rpseu-GMI1000) and reported a small number of putative false positives (22 and 1 genes, respectively). Nevertheless, our multi-replicate results are consistent with the targeted mutant validations reported by Su et al. (2021). Given our stringent filters and the variability for certain loci, our study may have failed to detect 11 additional genes that promote growth of Rpseu-GMI1000; we did not identify putative false positives in our dataset. These observations highlight the importance of biological replication for robust inference in RB-TnSeq studies.

By placing quantitative *in planta* fitness data in an evolutionary context, this work shows how ancestral physiological traits and lineage-specific innovations jointly shape the ability of *Ralstonia* wilt pathogens to colonize plant xylem. We reveal antimicrobial conflict as a recurring, evolutionarily selected feature of these pathogens’ life inside plant hosts. Together, these findings underscore the value of comparative functional genomics for understanding plant pathogenesis.

## Supporting Information

Supplementary figures: S1-S16

Supplementary tables: S1-S5

## Acknowledgements

Support was provided by the joint NSF / U.S. Department of Agriculture NIFA Plant Biotic Interactions program (NSF award #2336557, NIFA award #2024-67013-43303, and NIFA award #2023-67013-40245), the USDA Hatch Program (Project #1023861), and a Hellman Fellowship Foundation (to T. Lowe-Power). Nathalie Aoun was supported by USDA NIFA Fellowship #2024-67012-42635.

We thank the staff at the UC Berkeley CNR Oxford Tract Greenhouse and the UC Davis Controlled Environment Facility for maintaining the facilities where plants were grown. We thank Tyler Helmann and Steve Lindow for discussion of experimental design. We thank Prof. Daniel Runcie (Department of Plant Sciences, University of California, Davis) for helpful discussions on data analysis. We thank Ariana Enriquez for creating the cartoon illustrations of the flask and plant used in Figures 1 and S1.

This manuscript was edited with assistance from Microsoft Copilot. ChatGPT was used in to assist with troubleshooting R scripts.

## Author contributions

NA: Conceptualization, Data curation, Analysis, Funding acquisition, Experiments / Investigation, Methodology, Project administration, Resources, Software, Validation, Visualization, Writing – original draft, Writing – review & editing; SJG: Analysis, Writing – review & editing; AD: Analysis, Software, Writing – review & editing; TML: Conceptualization, Data curation, Analysis, Funding acquisition, Experiments / Investigation, Methodology, Project administration, Resources, Supervision, Validation, Visualization, Writing – original draft, Writing – review & editing

## Data availability

The BLASTp result datasets used to generate most figures are deposited in FigShare and are available at: 10.6084/m9.figshare.31796191.

---

**Figure S1.**
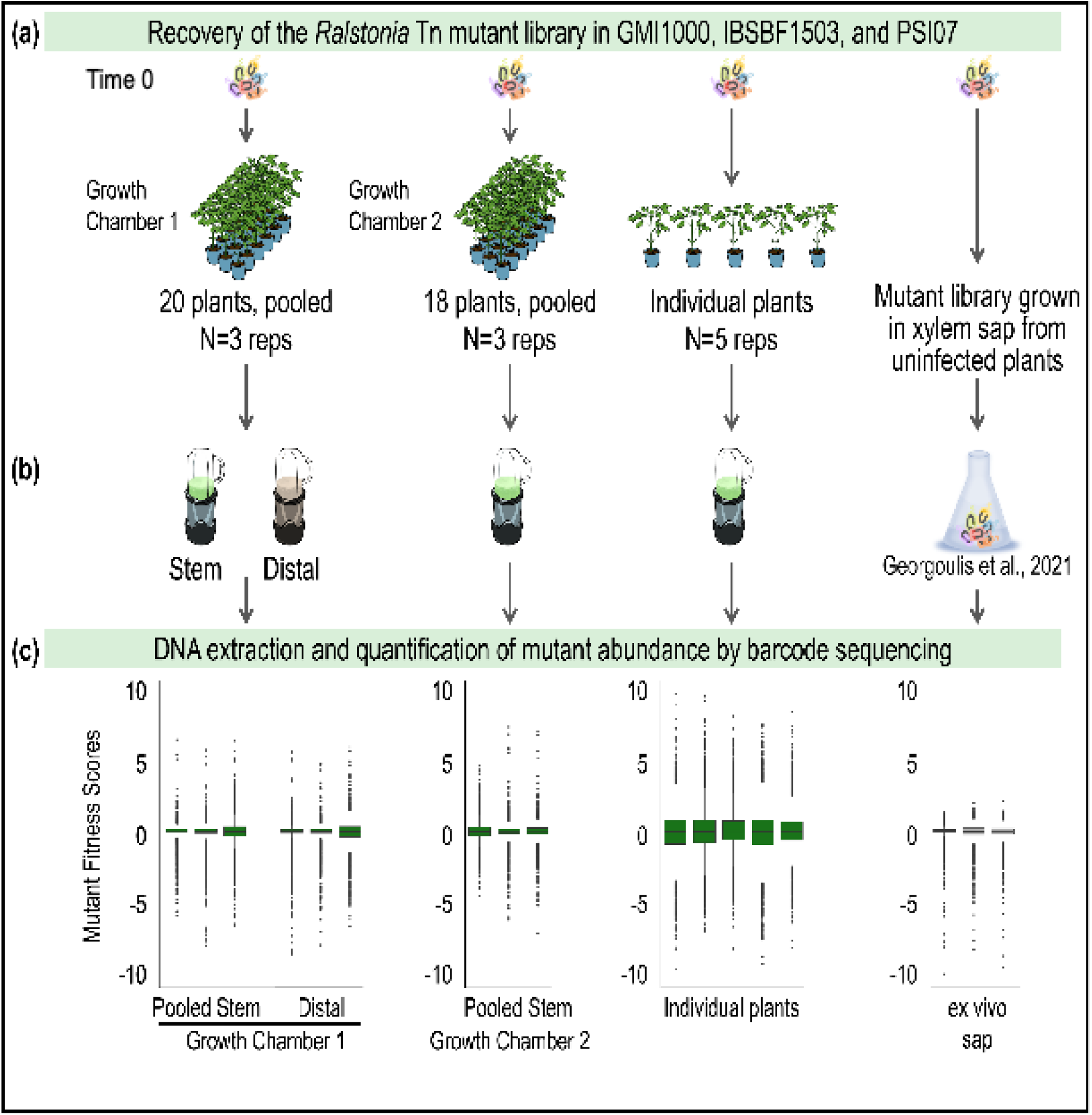
Comparisons of experimental workflows for RB-TnSeq experiments. (**A**) An RB-TnSeq experiment consists of growing the barcoded transposon mutant library in rich media and harvesting bacterial mutants from this inoculum (Time 0 control). The inoculum was used to inoculate the petiole of tomato *cv.* Moneymaker plants (N=20 and N=18) or filter-sterilized xylem sap harvested from healthy tomato plants [23]. (**B**) After disease onset, the *Ralstonia* transposon (Tn) mutant library was harvested after homogenizing pooled stems of the 20 or 18 plants, or from individual plants. Bacterial DNA was then extracted from Time 0 and experimental samples for amplicon barcode sequencing (BarSeq). (**C**) Fitness scores were calculated from BarSeq data as the log2 ratio of each mutant’s relative competitive success in the population compared with the initial inoculum. Gene fitness scores thus reflect the contribution of each gene to bacterial fitness under each condition. Experiments were performed using mutant libraries in three *Ralstonia* species complex backgrounds: Rpseu-GMI1000, Rsol-IBSBF1503, and Rsyz-PSI07.

**Figure S2.**
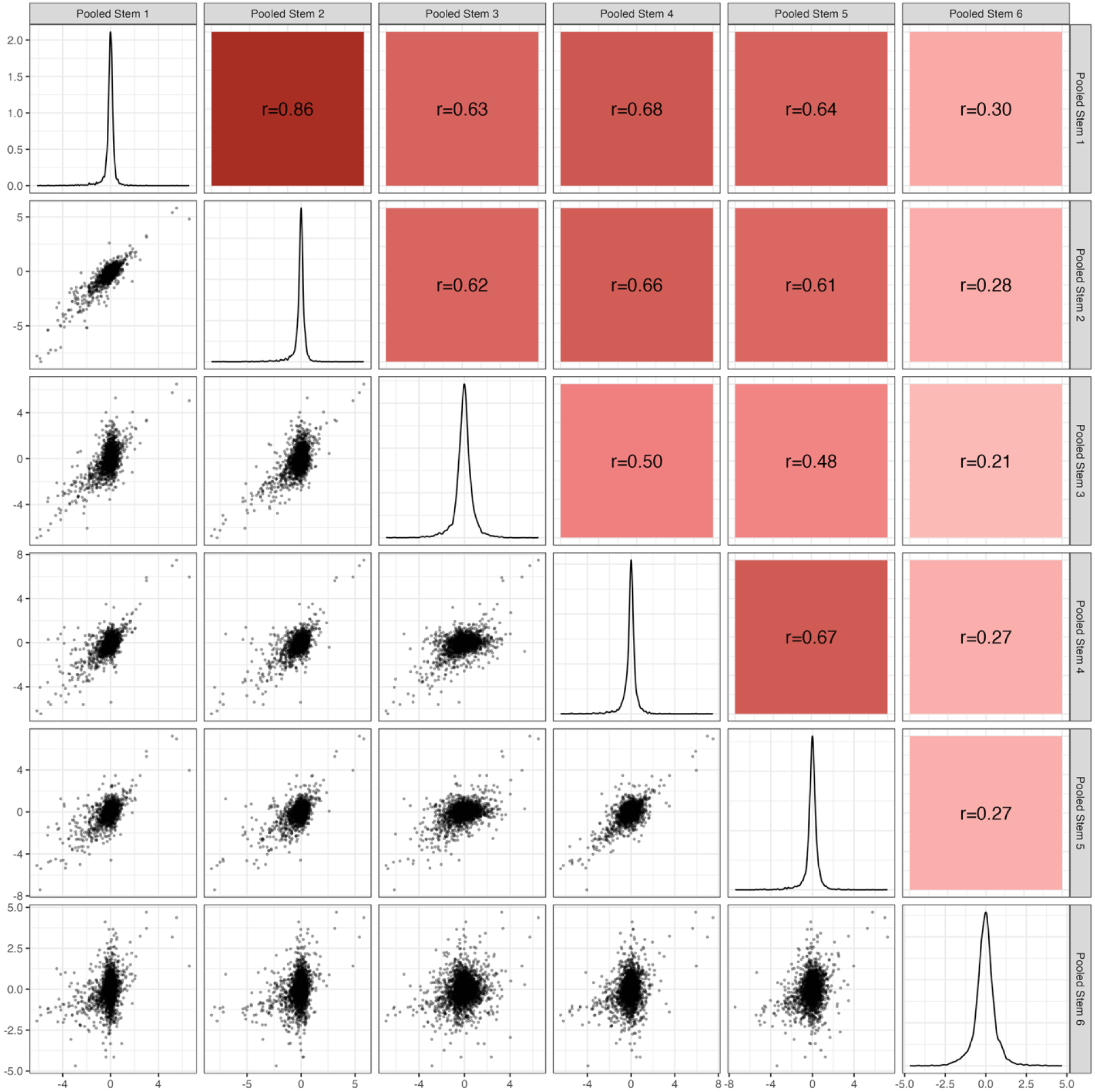
Pairwise correlations among six biological replicates of RB-TnSeq experiments in the pooled stem condition in the Rpseu-GMI1000 strain background. Pairwise scatterplots, density distributions, and Pearson correlation coefficients are shown for six pooled stem replicates (Pooled Stem 1-6). Diagonal panels display kernel density estimates for each pooled stem replicate. Lower panels show pairwise scatterplots with individual data points, illustrating the distribution and linear association between replicates. Upper panels report Pearson’s correlation coefficients (r), with background shading indicating correlation strength (darker red denotes stronger positive correlations). Based on low r, Rpseu-GMI1000 Pooled Stem replicate 6 was excluded from further analysis

**Figure S3.**
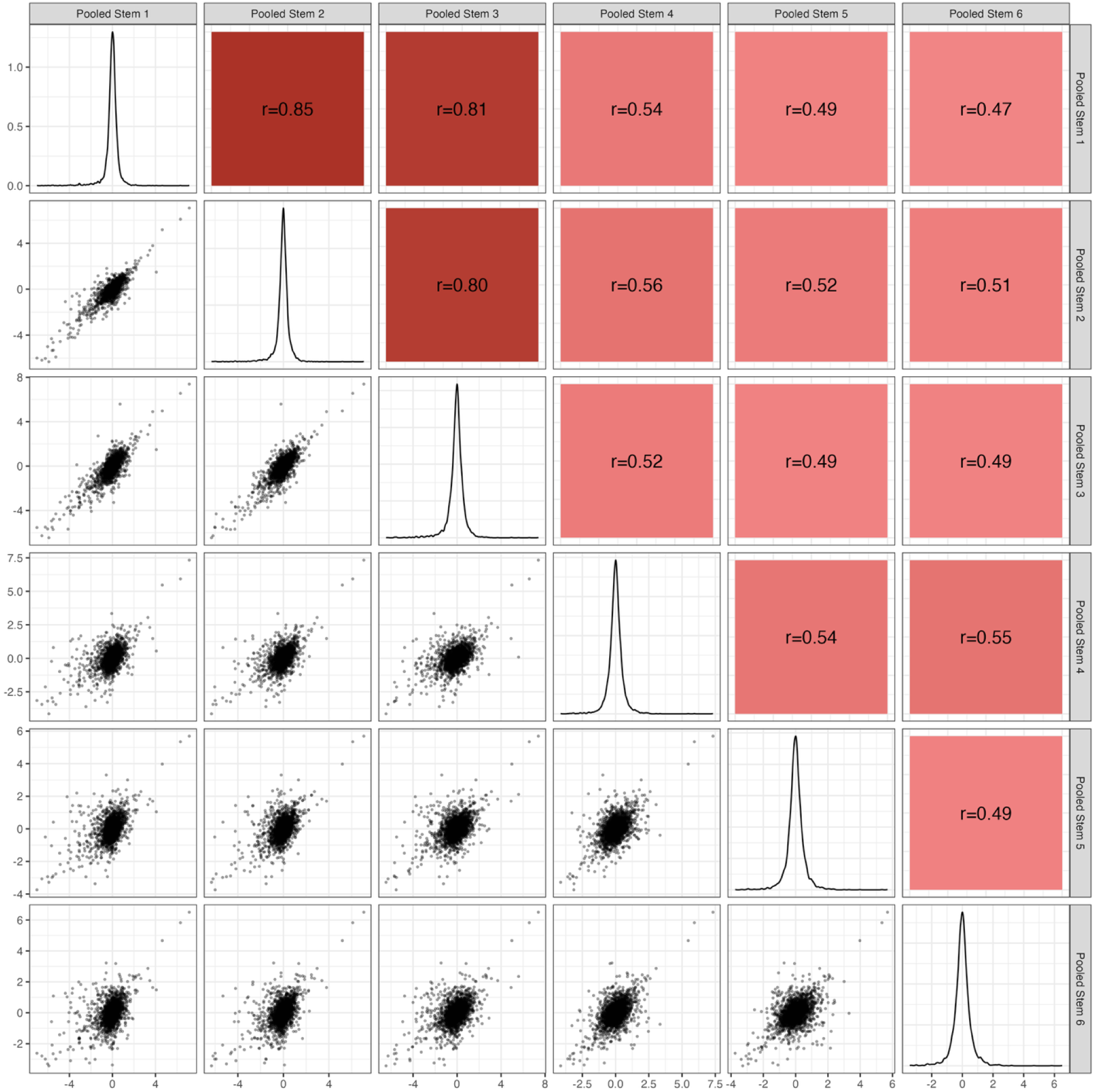
Pairwise correlations among biological replicates of RB-TnSeq experiments in the pooled stem condition in the Rsol-IBSBF1503 strain background. Pairwise scatterplots, density distributions, and Pearson correlation coefficients are shown for six pooled stem replicates (Pooled Stem 1-6). Diagonal panels display kernel density estimates for each pooled stem replicate. Lower panels show pairwise scatterplots with individual data points, illustrating the distribution and linear association between replicates. Upper panels report Pearson’s correlation coefficients (r), with background shading indicating correlation strength (darker red denotes stronger positive correlations).

**Figure S4.**
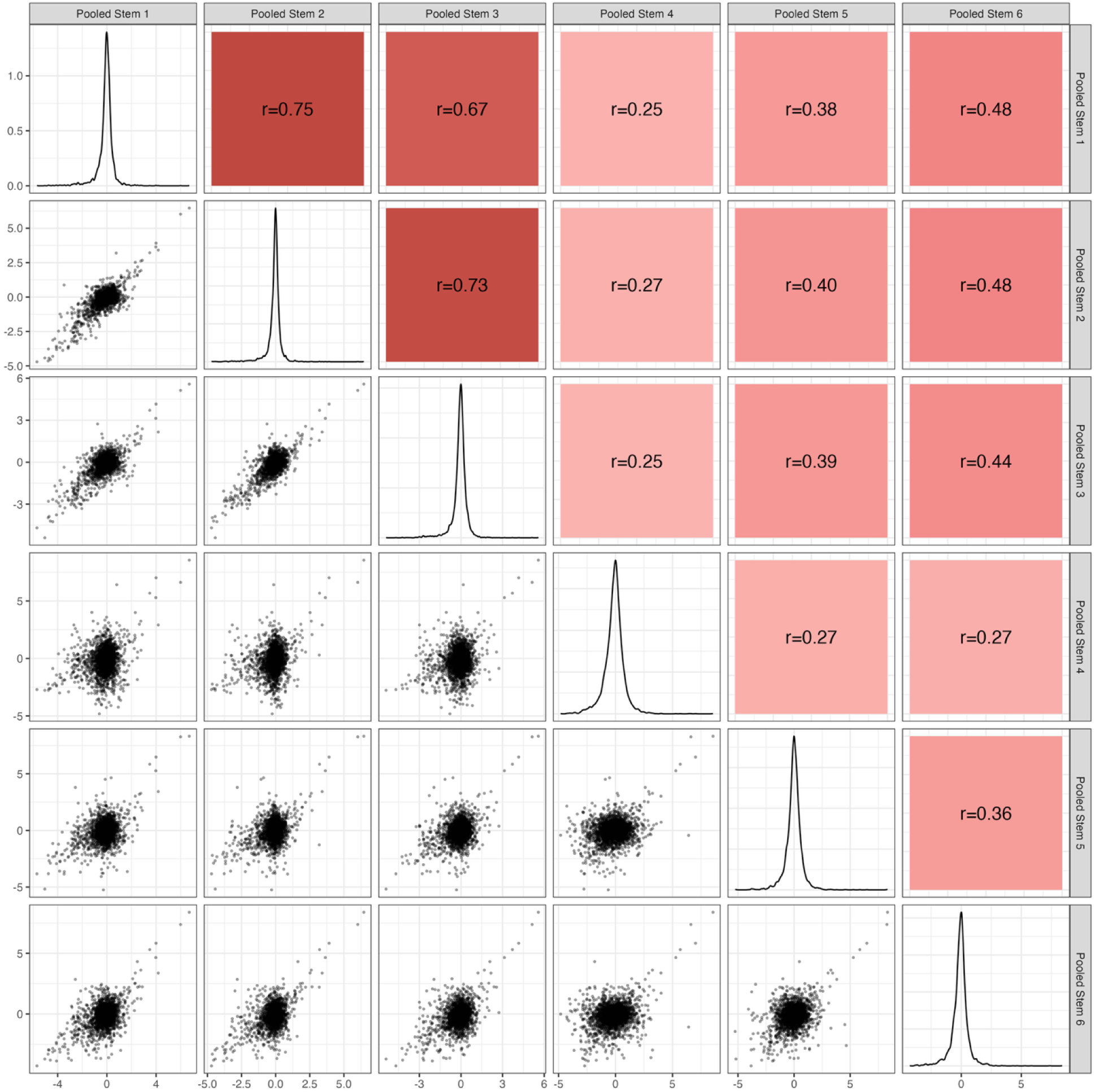
Pairwise correlations among biological replicates of RB-TnSeq experiments in the pooled stem condition in the Rsyz-PSI07 strain background. Pairwise scatterplots, density distributions, and Pearson correlation coefficients are shown for six pooled stem replicates (Pooled Stem 1-6). Diagonal panels display kernel density estimates for each pooled stem replicate. Lower panels show pairwise scatterplots with individual data points, illustrating the distribution and linear association between replicates. Upper panels report Pearson’s correlation coefficients (r), with background shading indicating correlation strength (darker red denotes stronger positive correlations). Based on low r, Rsyz-PSI07 Pooled Stem replicate 4 was excluded from further analysis.

**Figure S5.**
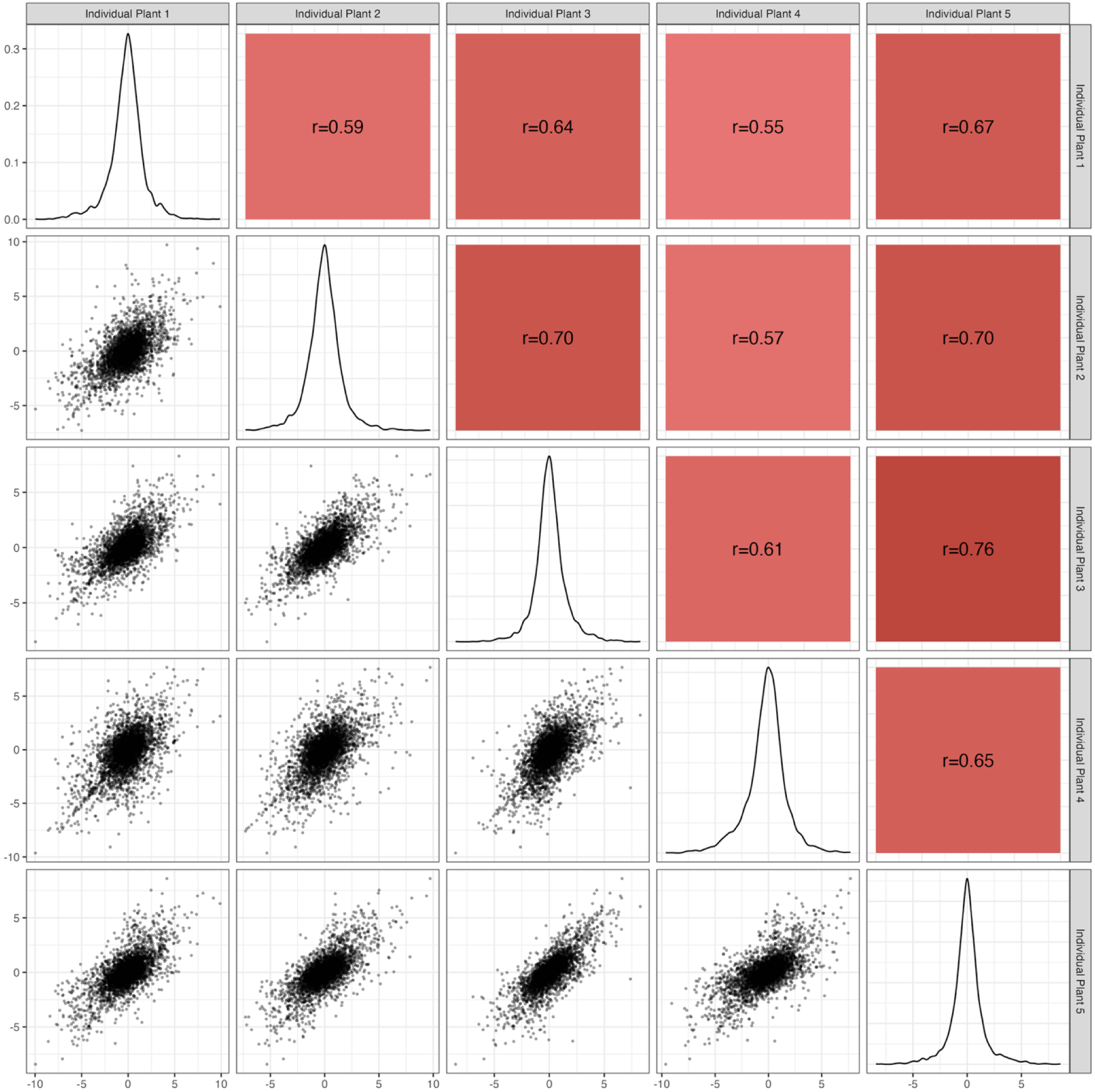
Pairwise correlations among five individual plants of RB-TnSeq in the Rpseu-GMI1000 strain background. Pairwise scatterplots, density distributions, and Pearson correlation coefficients are shown for five individual plants. Diagonal panels display kernel density estimates for each individual plant. Lower panels show pairwise scatterplots with individual data points, illustrating the distribution and linear association between replicates. Upper panels report Pearson’s correlation coefficients (r), with background shading indicating correlation strength (darker red denotes stronger positive correlations).

**Figure S6.**
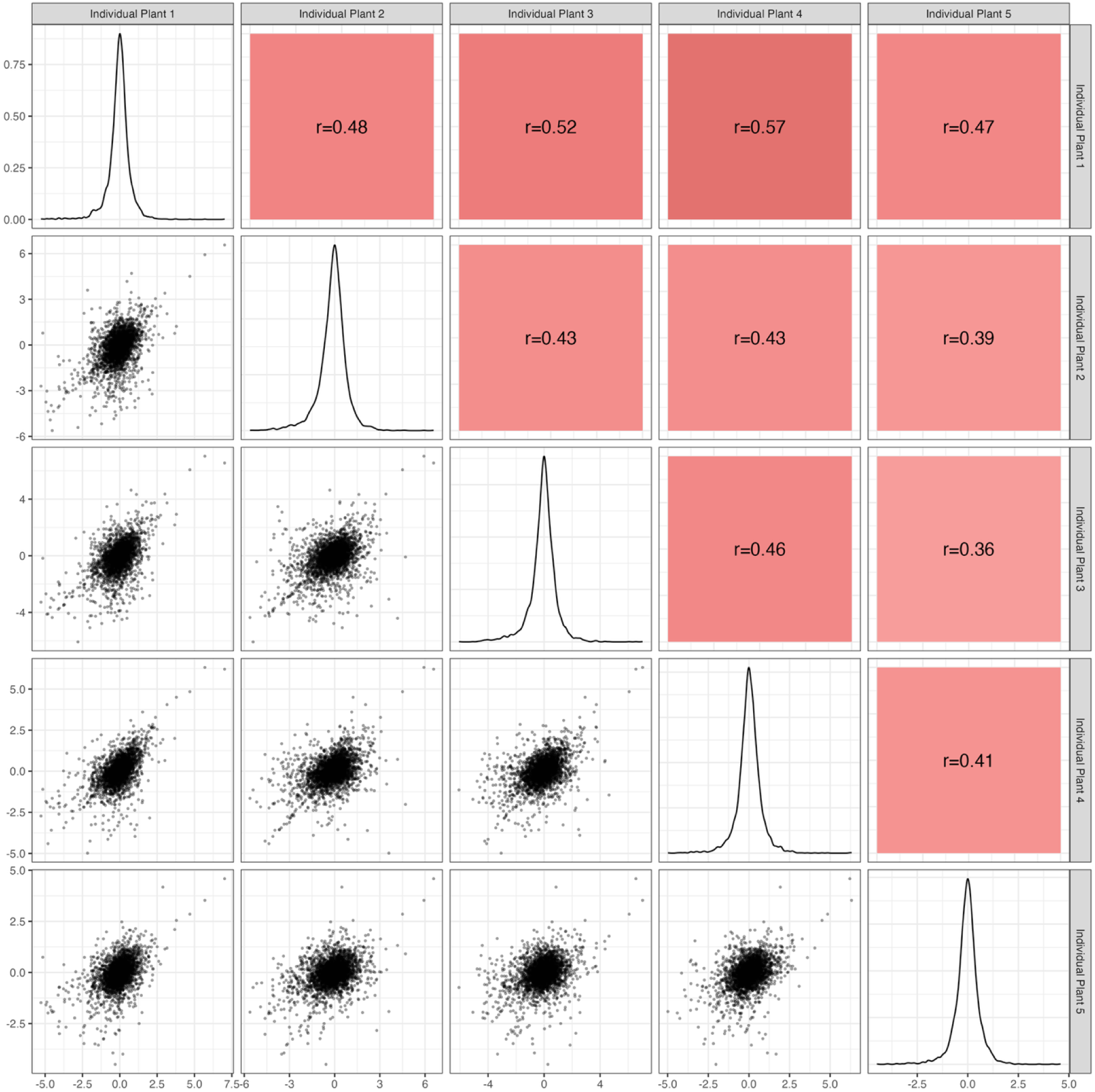
Pairwise correlations among five individual plants of RB-TnSeq in the Rsol-IBSBF1503 strain background. Pairwise scatterplots, density distributions, and Pearson correlation coefficients are shown for five individual plants. Diagonal panels display kernel density estimates for each individual plant. Lower panels show pairwise scatterplots with individual data points, illustrating the distribution and linear association between replicates. Upper panels report Pearson’s correlation coefficients (r), with background shading indicating correlation strength (darker red denotes stronger positive correlations).

**Figure S7.**
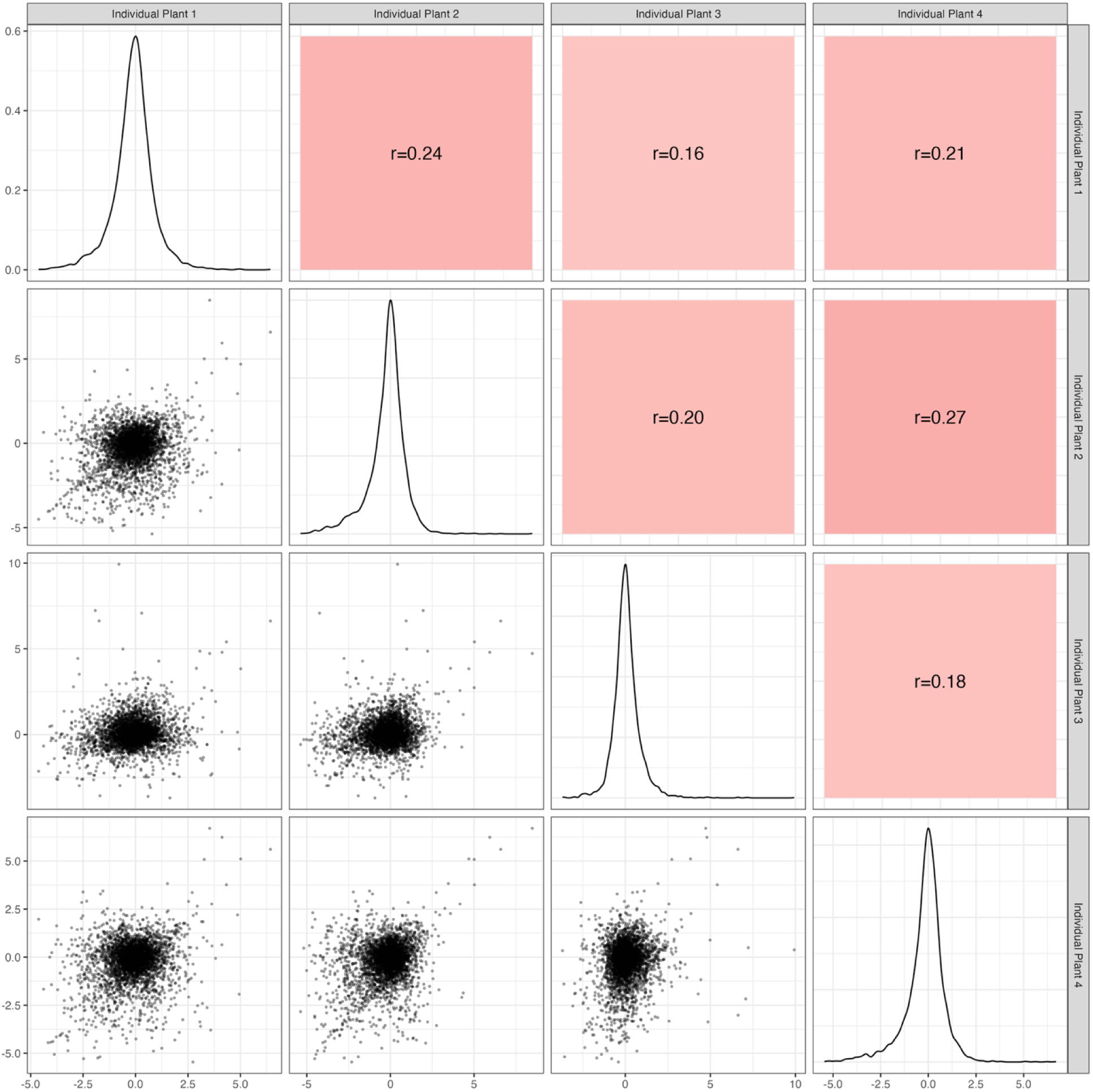
Pairwise correlations among four individual plants of RB-TnSeq in the Rsyz-PSI07 strain background. Pairwise scatterplots, density distributions, and Pearson correlation coefficients are shown for five individual plants. Diagonal panels display kernel density estimates for each individual plant. Lower panels show pairwise scatterplots with individual data points, illustrating the distribution and linear association between replicates. Upper panels report Pearson’s correlation coefficients (r), with background shading indicating correlation strength (darker red denotes stronger positive correlations).

**Figure S8.**
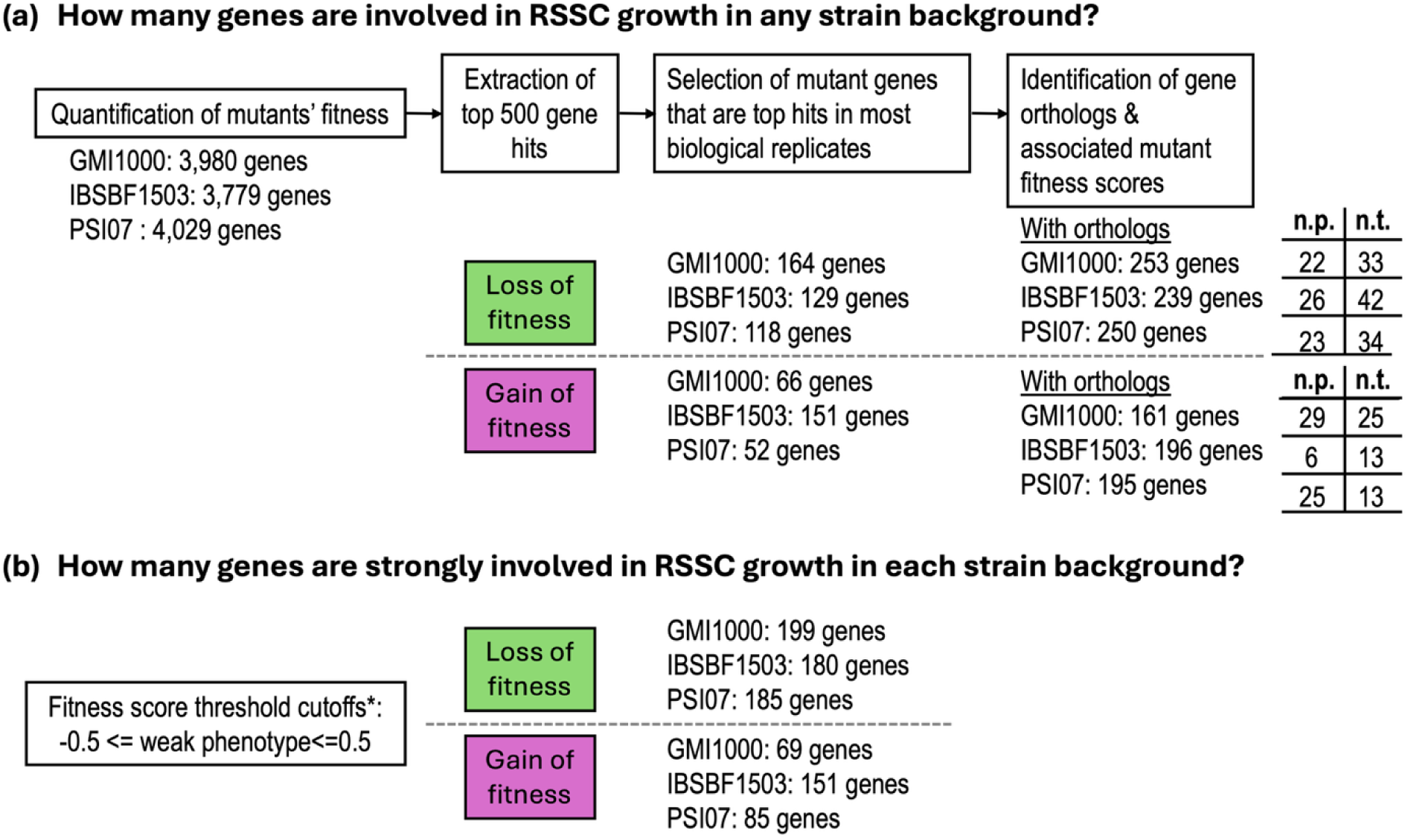
Workflow for identifying fitness-associated genes from pooled *in planta* RB-TnSeq experiments. (**a**) Mutant fitness was quantified under pooled *in planta* conditions, and the top 500 genes with the strongest fitness effects were extracted. Mutant genes were retained if they ranked among the top hits in at least 4 out of 5 or of 5 out of 6 biological replicates (see materials and methods). Syntenic orthologs across Rpseu-GMI1000, Rsol-IBSBF1503, and Rsyz-PSI07 were identified via pangenome analysis in KBase [34] and synteny analysis with Clinker [35]. (**b**) Gene-associated mutant fitness scores were classified as strong growth-promoting or -hindering genes based on fitness thresholds (fitness score≤“0.5 or ≥0.5). n.p.” indicates not present due to missing ortholog. “n.t. ” indicates not tested due to absence of the corresponding mutant in the RB-TnSeq library.

**Figure S9.**
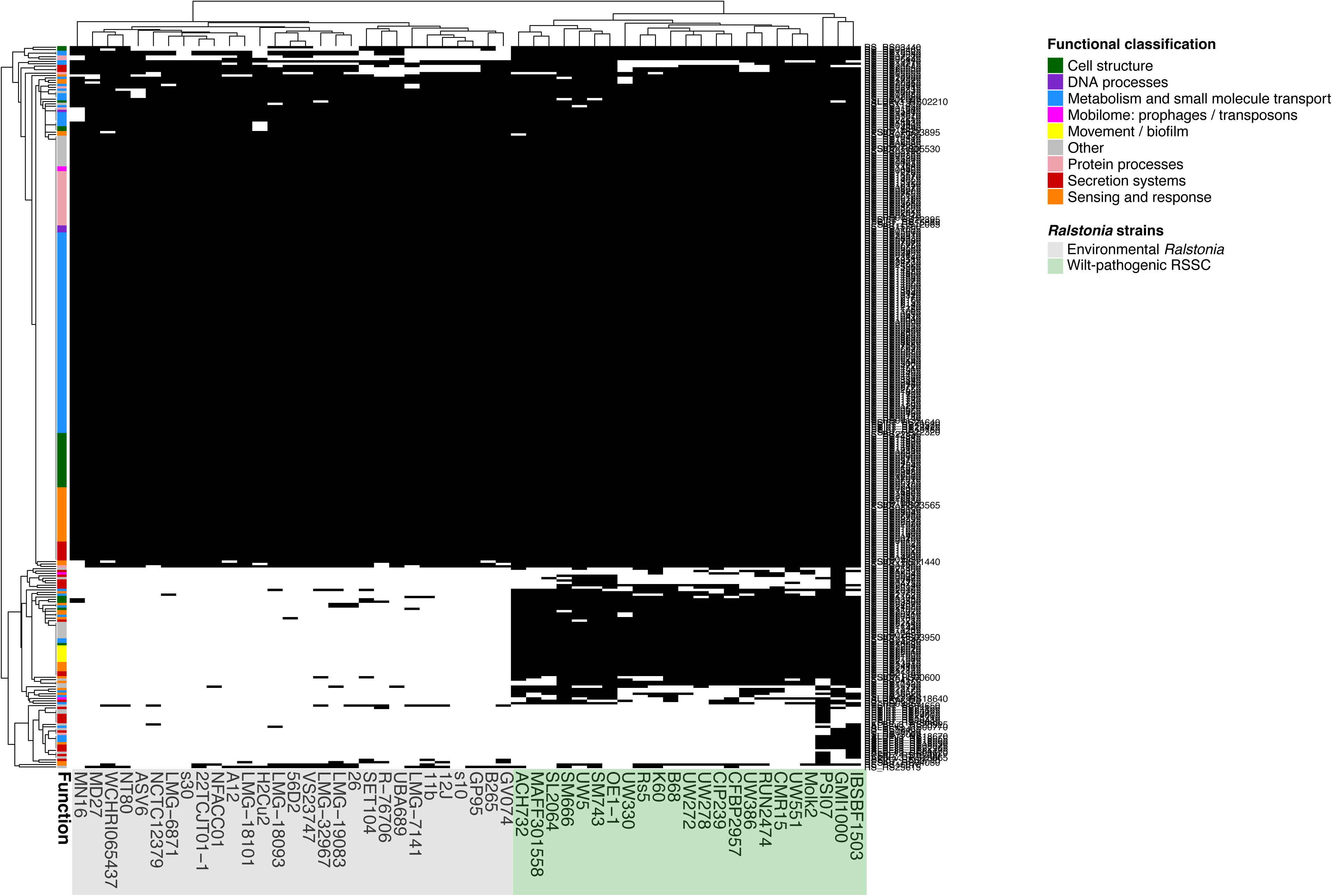
Presence-absence patterns of fitness-promoting genes distinguish wilt-pathogenic RSSC and environmental *Ralstonia* strains. Hierarchical clustering of *Ralstonia* strains based on gene abundance profiles reveals a clear separation between wilt-pathogenic RSSC and environmental *Ralstonia* isolates. Rows correspond to genes and columns to strains; black and white cells indicate gene presence and absence, respectively. Dendrograms were generated using hierarchical clustering based on gene abundance similarity, grouping wilt-pathogenic RSSC strains together and distinct from environmental *Ralstonia* strains. The colored bar alongside genes denotes functional classification.

**Figure S10.**
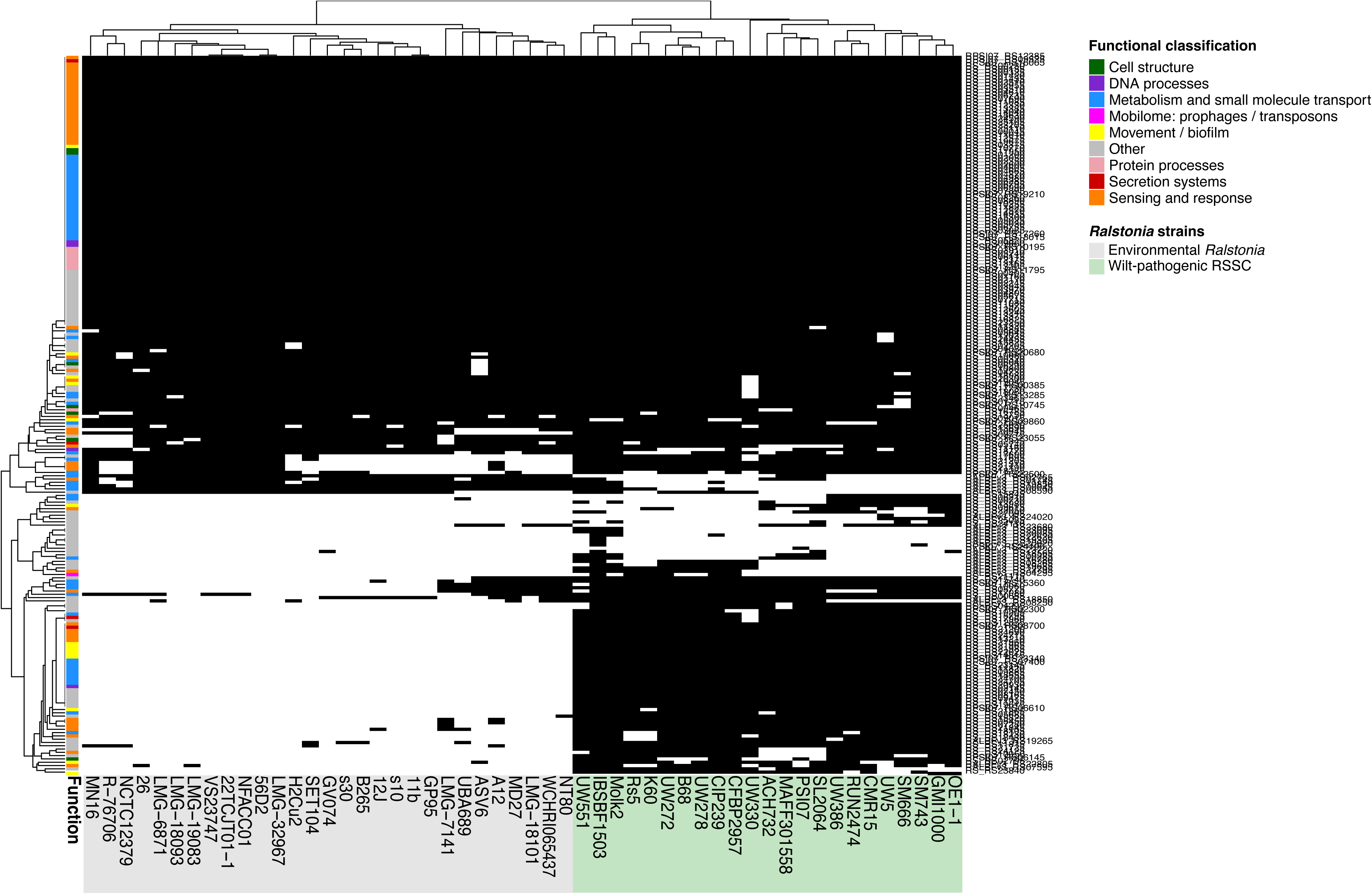
Presence-absence patterns of fitness-hindering genes distinguish wilt-pathogenic RSSC and environmental *Ralstonia* strains. Hierarchical clustering of *Ralstonia* strains based on gene abundance profiles reveals a clear separation between wilt-pathogenic RSSC and environmental *Ralstonia* isolates. Rows correspond to genes and columns to strains; black and white cells indicate gene presence and absence, respectively. Dendrograms were generated using hierarchical clustering based on gene abundance similarity, grouping RSSC strains together and distinct from environmental strains. The colored bar alongside genes denotes functional classification.

**Figure S11.**
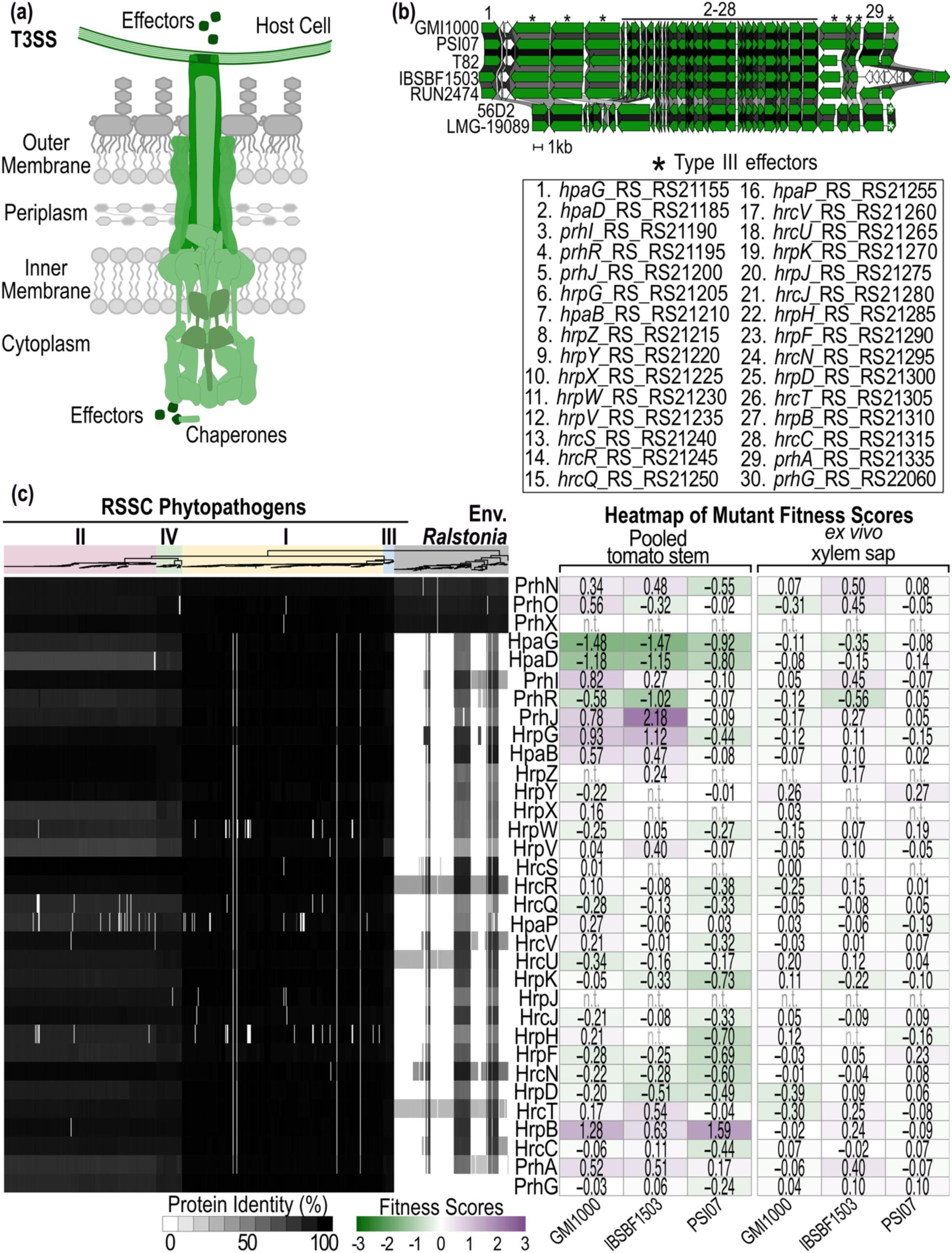
Conversation and fitness impacts of the T3SS gene cluster. (**a**) Schematic representation of T3SS core structural components in *Ralstonia*. (**b**) Genetic organization of the T3SS in RSSC (Rpseu-GMI10000, Rsys-PSI07, Rsyz-T82, Rsol-IBSBF1503, and Rpseu-RUN2474) compared to environmental isolates (*R. wenshanensis* 56D2 and *Ralstonia edaphi* LMG-19089). (**c**) Conservation of T3SS proteins across the *Ralstonia* genus (n=536 plant-pathogenic RSSC genomes and n=150 other *Ralstonia* genomes). Proteins from Rpseu-GMI1000 were queried and sequence identity of the best BLASTp hit with ≥60% coverage is shown as greyscale heatmaps below the phylogenetic tree. Due to a systematic annotation bias in which the Prokka implementation in KBase fails to annotate intact *prhA* genes in RSSC phylotype II genomes, 85 phylotype II genomes were excluded; the retained 202 phylotype II genomes are RefSeq assemblies annotated using NCBI PGAP. *In planta* and xylem sap fitness score of homologs in RSSC species representatives (Rpseu-GMI1000, Rsol-IBSBF1503, and Rsyz-PSI07) are shown as green-to-purple heatmaps on the right. “n.t.” indicates not tested due to absence of the corresponding mutant in the RB-TnSeq library. The phylogenetic tree was generated in KBase using FastTree2 from a multiple sequence alignment (MSA) of 49 conserved bacterial genes [41]. Colored backgrounds denote phylotypes of plant pathogenic RSSC versus environmental *Ralstonia* strains. The phylogenetic tree and BLASTp results were visualized using iTOL [42], and fitness heatmaps were generated using ‘pheatmap’ in RStudio. The Figure was refined in Affinity Designer.

**Figure S12.**
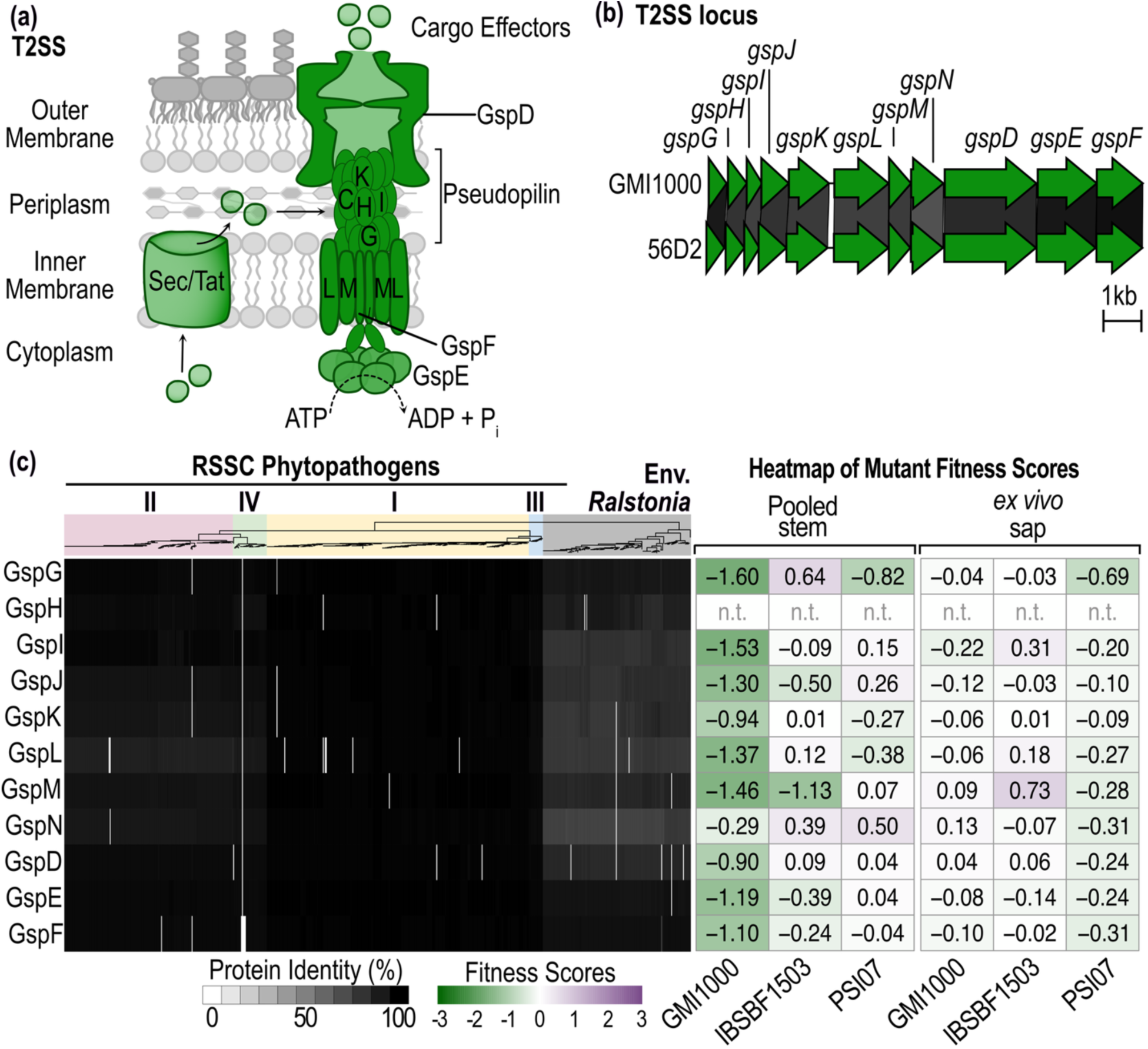
The broadly conserved type II secretion system (T2SS) contributes to Rpseu*-*GMI1000 fitness *in planta*. (**a**) Schematic representation of the T2SS. (**b**) Genetic organization of the T2SS in Rpseu-GMI1000 strain (RSSC) and the environmental *R. wenshanensis* 56D2 strain. (**c**) Distribution of T2SS structural proteins (GspG-GspF) based on percent protein identity across 505 plant-pathogenic RSSC strains and 150 environmental *Ralstonia* strains. Proteins from Rpseu-GMI1000 were queried and sequence identity of the best BLASTp hit with ≥60% coverage is shown as greyscale heatmaps below the phylogenetic tree. Due to a systematic annotation bias where the Prokka implementation in KBase fails to annotate intact *gspD* genes in RSSC phylotype II genomes, 116 phylotype II genomes were excluded from this figure; the retained 171 phylotype II genomes are RefSeq assemblies annotated using NCBI PGAP. *In planta* and xylem sap fitness score of homologs in RSSC species representatives (Rpseu-GMI1000, Rsol-IBSBF1503, and Rsyz-PSI07) are shown as green-to-purple heatmaps on the right. “n.t. ” indicates not tested due to absence of the corresponding mutant in the RB-TnSeq library. The phylogenetic tree was generated in KBase using FastTree2 from a multiple sequence alignment (MSA) of 49 conserved bacterial genes [41]. Colored backgrounds denote phylotypes of plant pathogenic RSSC versus environmental *Ralstonia* strains. The phylogenetic tree and BLASTp results were visualized using iTOL [42], and fitness heatmaps were generated using ‘pheatmap’ in RStudio. The Figure was refined in Affinity Designer.

**Figure S13.**
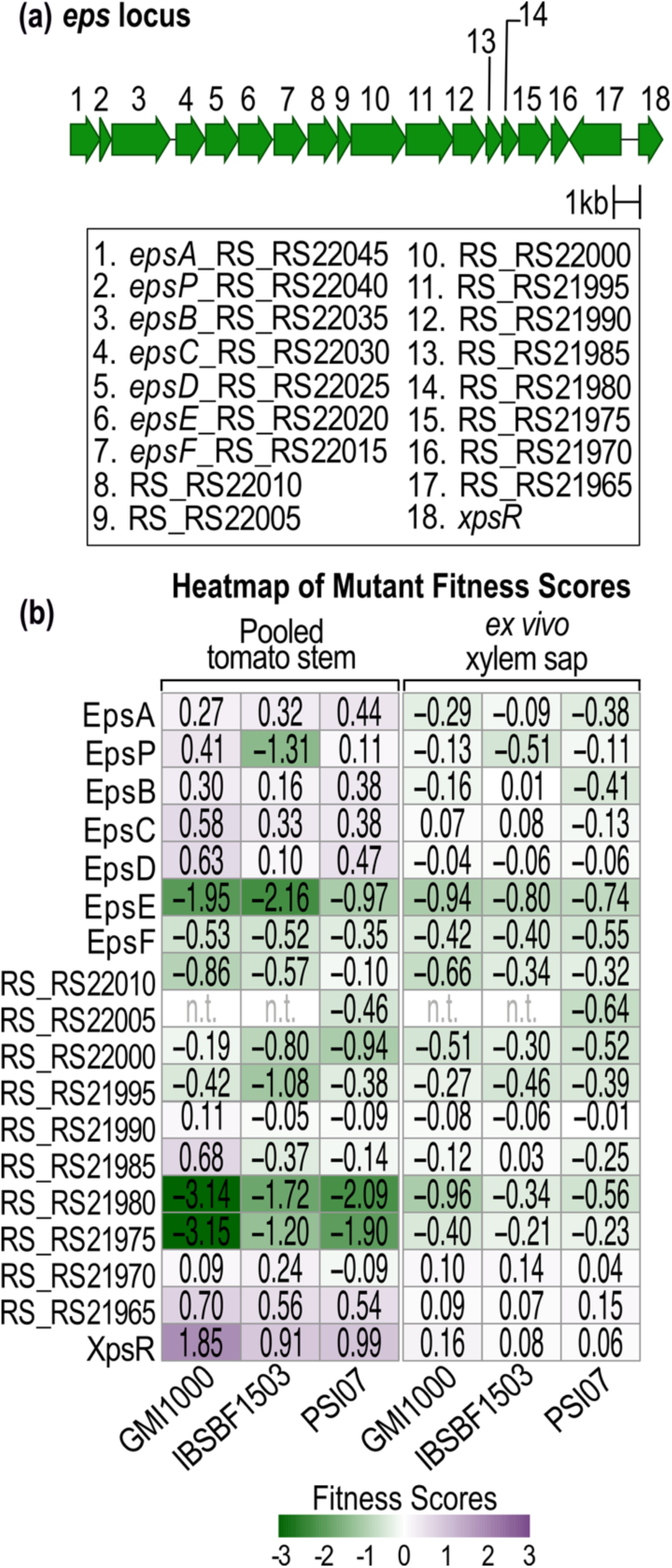
Fitness patterns of EPS-I biosynthesis genes are complex. (**a**) Organization of the *eps* gene cluster. (**b**) Heatmaps of mutant fitness scores across Rpseu-GMI1000, Rsol-IBSBF1503, and Rsyz-PSI07 under pooled tomato stem and xylem sap conditions. *In planta* and xylem sap fitness score of homologs in RSSC species representatives (Rpseu-GMI1000, Rsol-IBSBF1503, and Rsyz-PSI07) are shown as green-to-purple heatmaps on the right. “n.t. ” indicates not tested due to absence of the corresponding mutant in the RB-TnSeq library. Fitness heatmaps were generated using ‘pheatmap’ in RStudio. The Figure was refined in Affinity Designer.

**Figure S14.**
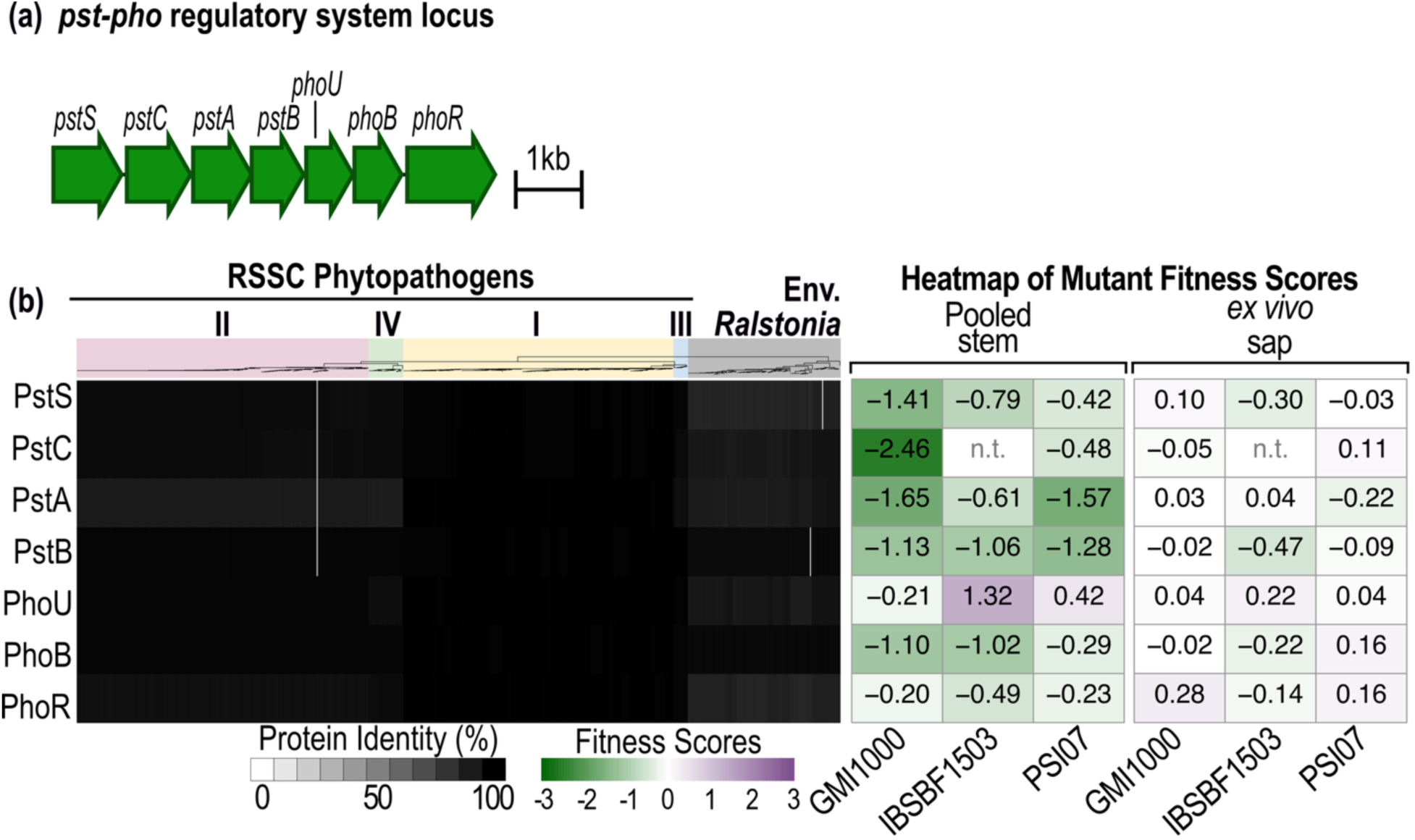
Genes in the highly conserved Pst transporter and Pho regulatory pathways are required for RSSC growth *in planta*. (**a**) Genetic organization of *pstSCAB* and *phoUBR*. (**b**) Phylogenetic distribution of Pst and Pho proteins based on percent protein identity across 621 plant-pathogenic RSSC and 150 environmental *Ralstonia* strains. Proteins from Rpseu-GMI1000 were queried and sequence identity of the best BLASTp hit with ≥60% coverage is shown as greyscale heatmaps below the phylogenetic tree. Heatmaps on the right show RB-TnSeq fitness scores for the genes in Rpseu-GMI1000, Rsol-IBSBF1503, and Rsyz-PSI07 backgrounds under pooled tomato stem and xylem sap conditions. “n.t. ” indicates not tested due to absence of the corresponding mutant in the RB-TnSeq library. The phylogenetic tree was generated in KBase using FastTree2 from a multiple sequence alignment (MSA) of 49 conserved bacterial genes [41]. Colored backgrounds denote phylotypes of plant pathogenic RSSC versus environmental *Ralstonia* strains. The phylogenetic tree and BLASTp results were visualized using iTOL [42], and fitness heatmaps were generated using ‘pheatmap’ in RStudio. The Figure was refined in Affinity Designer.

**Figure S15.**
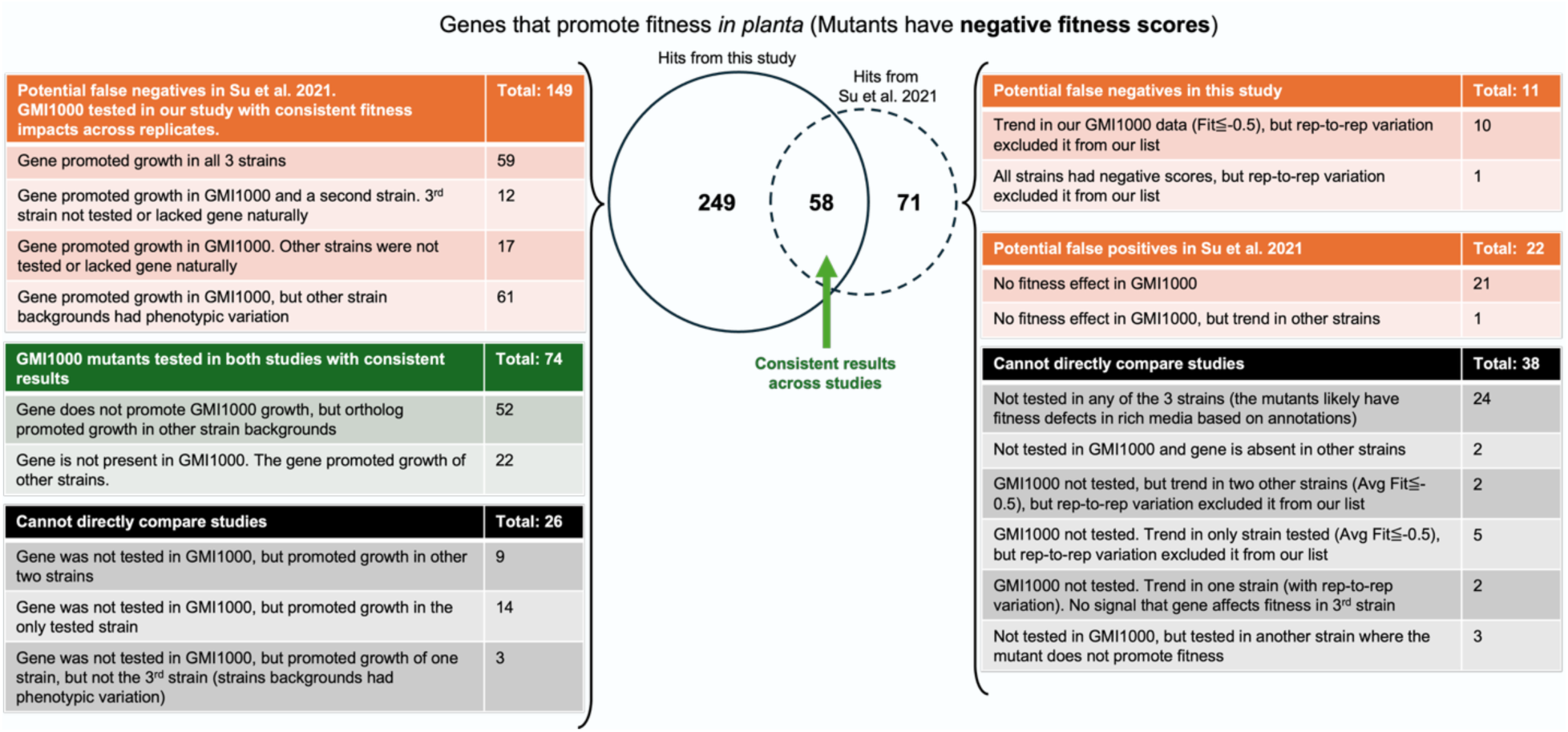
Comparison of *in planta* RB-TnSeq growth-promoting fitness genes between this study and Su et al. (2021). Fitness effects of mutant genes identified in this study were compared with those reported by Su *et al.,* 2021 [18]. The overlapping Venn diagram highlights growth-promoting genes identified in both studies. The left side of the Venn diagram (N = 249 genes) includes 149 potential false negatives in Su *et al.* (2021) based on consistent measurements in our replicated dataset, 74 where the Rpseu-GMI1000 mutants had consistent phenotypes in both studies, and 26 factors where direct comparisons across studies were not possible. On the right side of the Venn diagram (N = 71 genes), counts refer to 11 genes representing potential false negatives in our study, 22 potential false positives reported by Su *et al.* (2021), and 38 factors where direct comparisons across studies were not possible.

**Figure S16.**
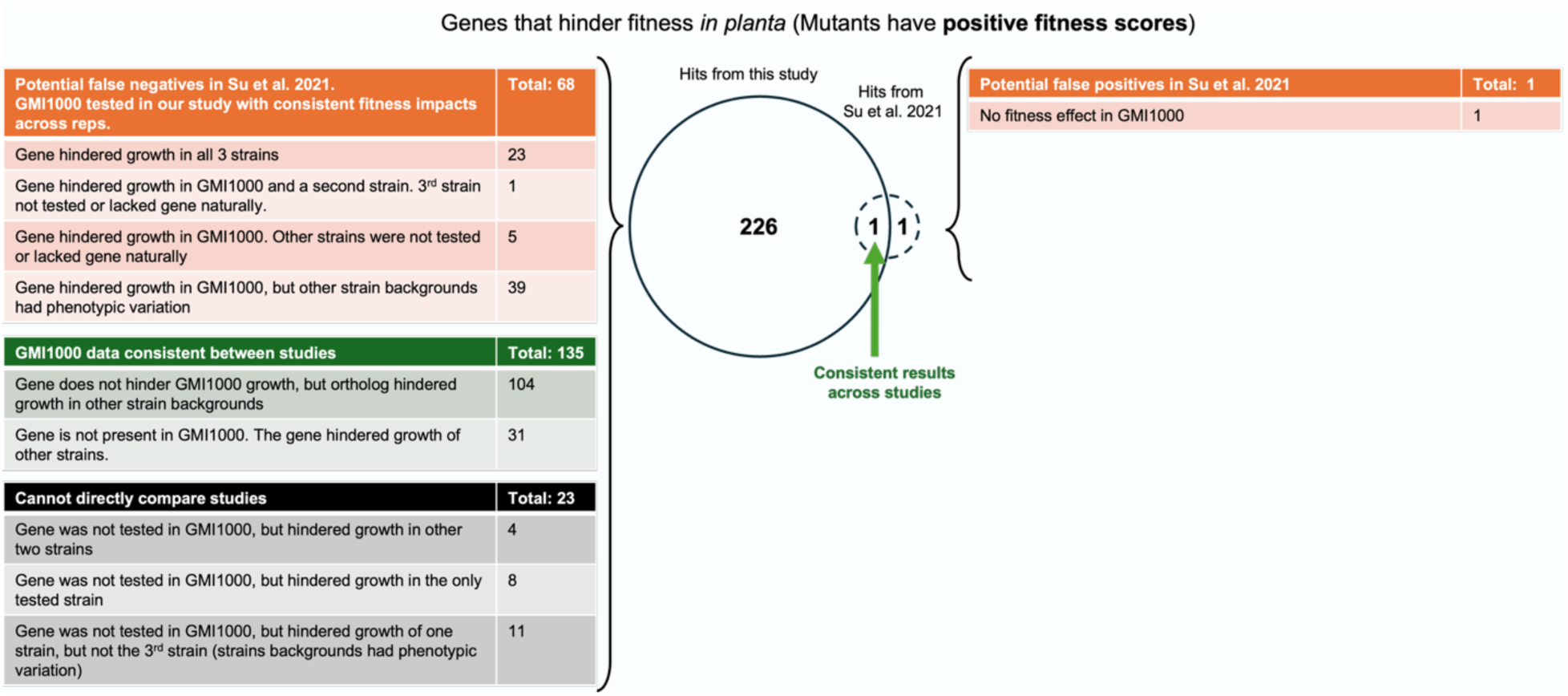
Comparison of *in planta* RB-TnSeq growth-hindering fitness genes between this study and Su et al. (2021). Fitness effects of mutant genes identified in this study were compared with those reported by Su *et al.,* 2021 [18]. The overlapping Venn diagram highlights one growth-hindering gene, *efpR*, identified in both studies. The left side of the Venn diagram (N = 226 genes) includes 68 potential false negatives in Su *et al.,* 2021 based on consistent measurements in our replicated dataset, 135 where the Rpseu-GMI1000 mutants had consistent phenotypes in both studies, and 23 factors where direct comparisons across studies were not possible. On the right side of the Venn diagram is one potential false positive reported by Su *et al.,* 2021.

